# Disentangling the roles of inter- and intraspecific variation on leaf trait distributions across the eastern United States

**DOI:** 10.1101/2021.04.01.438064

**Authors:** Sergio Marconi, Benjamin G. Weinstein, Jeremy W. Lichstein, Stephanie A. Bohlman, Aditya Singh, Ethan P. White

## Abstract

Functional traits are influenced by phylogenetic constraints and environmental conditions, but previous large-scale studies modeled traits either as species weighted averages or directly from the environment, precluding analyses of the relative contributions of inter- and intraspecific variation across regions. We developed a joint model integrating phylogenetic and environmental information to understand and predict the distribution of eight leaf traits across the eastern USA. This model explained 68% of trait variation, outperforming both species-only and environment-only models, with variance attributable to species alone (23%), the environment alone (13%), and their combined effects (25%). The importance of the two drivers varied by trait. Predictions for the eastern USA produced accurate estimates of intraspecific variation and deviated from both species-only and environment-only models. Predictions revealed that intraspecific variation holds information across scales, affects relationships in the leaf economic spectrum and is key for interpreting trait distributions and ecosystem processes within and across ecoregions.

## Introduction

Global change is expected to cause extensive changes in ecosystems, driving unprecedented redistribution of species and altering their traits (Diaz and Cabido, 2001). Understanding how these changes in the environment will drive adaptations within and across species is essential for management and conservation. Functional traits are among the most important properties of biological organisms because they are involved in key ecosystem processes from local community assembly (McGill et al., 2006, Sterck et al., 2011) to global biogeochemical cycles (Bonan et al., 2002; Verheijen et al., 2013), and these processes are interconnected across scales (Reichstein et al., 2014, Peaucelle et al., 2019). Biotic interactions, micro-climate, and soil conditions can affect species co-occurrence and influence local trait distributions (Bruelheide et al., 2018, Simpson et al., 2016), and variation in climate within species ranges can affect realized niches and drive trait responses (Chave, 2013). Relationships between traits, like the the leaf economic spectrum (LES) in plants (Wright et al., 2004), reveal information about biological constraints in resource allocation than impact ecophysiology and are a key component of ecosystem models (Fisher et al, 2015). Given their central role across levels of organization, understanding how traits vary within and among species across scales and environments is essential for conserving ecosystem function (Violle et al., 2014).

Traits vary geographically through a combination of shifts in species abundances and intraspecific trait variation (Leps et al., 2011, Laughlin et al., 2012, Münzbergová et al., 2017). Thus, understanding how traits respond to the environment across wide geographic areas requires approaches that integrate both inter- and intraspecific variation. This is challenging because individual level trait data are geographically and taxonomically limited, making it hard for traditional methods to identify the relative importance of inter- and intraspecific variation at large scales (Henn et al., 2018). In addition, due to limited data availability, studying traits at broad geographic scales may require predicting trait values for species that have not been sampled throughout their geographic range, or even species that have not been sampled at all. Given these challenges, most current approaches to studying broad-scale trait variation focus on predicting community weighted mean (CWM) trait values, which circumvents data limitations by either focusing directly on trait–environment relationships (based on direct relationships between environment and CWMs without explicitly considering species distributions or abundances) or by estimating CWMs directly from species mean trait values (without accounting for intraspecific variation, and considering environmental variables only for the purpose of predicting species distributions) (Miller & Ives, 2019).

Models based on direct trait-environment relationships offer the advantage of predicting trait distributions without requiring data on species abundances. A corollary of this approach is that it ignores well-established species and phylogenetic signals in trait variation driven by biological, physical, and historical constraints (Wright et al., 2004, Ordoñez et al., 2009, Anderegg et al., 2018), and implicitly assumes that the environment captures relevant changes in species distribution and abundance. However, since species-environment relationships are typically noisy (Clark 2016), failing to explicitly account for species distribution and abundance inevitably introduces prediction errors. Furthermore, models that do not explicitly incorporate species effects cannot be used to study inter- and intraspecific contributions to trait variation, which has important implications for functional ecology, ecophysiology, and ecosystem modeling (Anderegg et al., 2018, Osnas et al., 2018, Tautenhahn et al., 2020).

An alternative to pure environmental models is to predict traits directly from species-level means, which may provide better estimates of large-scale trait distributions (Clark 2016, Wieczynski et al., 2019, Swenson & Weiser, 2010). Species-based approaches assume that environmental drivers affect trait distributions only indirectly by shaping community assembly and therefore that species distributions themselves are the best predictor of traits and associated ecosystem function. Species-based models can be used to make predictions for traits over large areas by leveraging national-scale forest inventories or other vegetation networks and can be combined with species distribution models to forecast shifts in trait distributions (Swenson & Weiser, 2010, Clark 2016). However, species-level methods ignore intraspecific variation, which can be larger than interspecific variation for broadly distributed species (Niinemets, 2015, Messier et al., 2017).

Both pure environment and pure species approaches have contributed to our understanding of trait variation and allow for predicting community level trait distributions without requiring extensive field surveys of traits. However, as noted above, neither approach is designed to study intraspecific trait variation: pure environment approaches cannot separate inter- and intraspecific variation, and pure species approaches completely ignore intraspecific variation. Also, these approaches often fail to account for phylogenetic signals among closely related species (but see Swenson et al., 2017), which is potentially important for generating more robust trait predictions for species that are sparsely sampled, or not sampled at all (Blomberg et al., 2003, Swenson 2013, Swenson et al., 2014). These limitations prevent assessment of the relative importance of inter- and intraspecific variation on trait distributions across a continuum of geographical scales, reducing our ability to generalize and understand mechanisms driving trait distributions and inter-trait relationships.

To address these limitations, we developed a model that combines species identity (including phylogenetic relationships from the Tree of Life; Hinchliff et al., 2015) and environmental drivers (climate and topography) with leaf trait data from the National Ecological Observatory Network (NEON) (NEON, 2020). Our approach makes it possible to estimate the relative contributions of inter- and intraspecific variation to trait distributions at any location or scale where species abundance data are available. This allows us to address whether changes in the environment have a direct effect on trait distributions and inter-trait relationships, within and across species. We jointly modeled eight leaf traits: nitrogen (N%), carbon (C%), chlorophyllA (ChlA%), chlorophyllB (ChlB%), carotenoids (Crt%), leaf mass per area (LMA, g m^-2^), lignin (%) and cellulose (%). We compared our combined species-environment model to models based on only environmental drivers or only on species and phylogenetic information. We integrated the combined model with Forest Inventory and Analysis (FIA) data (Bechtold & Patterson. 2005) and Daymet data (Thornton et al., 2018) to make leaf trait predictions for ∼1.2 million trees across the eastern USA and validated the predictions against independent field datasets.

We compared predictions from our combined model to predictions from environment-only and species-only approaches to assess the influence of inter-vs. intraspecific variation on trait distributions across scales, species, and functional types. Specifically, we present four applications, in which we: (1) quantify the relative contributions of environmental factors vs. species and phylogenetic information to leaf trait variation at the continental scale; (2) assess drivers of ecoregional differences in trait distributions; (3) analyze large-scale intraspecific trait variation driven by environmental drivers; and (4) explore how intraspecific trait variation impacts the relationship between N% and LMA (one of the LES relationships) and the implications for representing leaf trait variation in ecosystem models.

## Materials and Methods

### Data

We used data from NEON (National Ecological Observatory Network, 2020), the Botanical Information and Ecology Network (BIEN, Maitner et al., 2020) and TRY (Kattge et al., 2020) to link information on leaf traits, species identity, and approximate locations for individual trees. We used Foliar Physical and Chemical Properties (DP1.10026.001) and Vegetation Structure data (DP1.10098.001) from NEON to build joint trait distribution models with environmental drivers (climate and topography) alone, phylogenetic drivers (species identity and phylogeny) alone, and both (combined model). Linking the two different NEON datasets produced individual tree data with stem geolocation and measures of eight leaf traits (LMA, chlorophyll A and B, carotenoids, lignin, cellulose, C, N) for 477 trees in 20 sites across the USA (Fig. S.1). Since foliar trait concentrations vary significantly with phenology and canopy position (Niinemets et al., 2015), foliar samples were collected at the “peak of greenness” and from the sunlit portion of the canopy. We tested the generalizability of our approach outside of NEON by evaluating predictions from independent (out-of-sample) eastern USA data available from the BIEN and TRY datasets (Appendix S1). These two datasets provide measures for C, N and LMA for a total of 353 individual trees. We used data from the Open Tree of Life (Redelings 2017) to measure phylogenetic distances among species, which we used to inform the covariance structure of our combined and species-only models.

Data for environmental drivers included average monthly climate data from 1995 to 2015 (Appendix S1) extracted from Daymet (Thornton et al., 2018) and topographic variables (elevation, slope, and aspect) reported in the NEON and FIA datasets. For three common eastern USA tree species (*Acer rubrum, Fagus grandifolia*, and *Abies balsamea*), we used all publicly available leaf N% data from the TRY database to quantify intraspecific variation in leaf N% across each species’ geographic range in the USA. We selected these three species because: (1) *Abies balsamea* is the needleleaf species with the most leaf N% data in TRY in the USA; (2) *Fagus grandifolia* is the broadleaf species with the most leaf N% data in TRY in the USA; and (3) *Acer rubrum* occurs throughout much of the eastern USA in a wide variety of habitats (e.g., from xeric to mesic) and has abundant leaf N% data in TRY. We combined our trait modeling approach with data from the USA Forest Inventory and Analysis (https://www.fia.fs.usda.gov/) (FIA) to estimate traits for all trees surveyed in the FIA across the eastern USA from 2016 to 2019. We used Lv.3 ecoregions and Lv.2 ecoprovinces as defined by the Environmental Protection Agency (Omernik et al., 2014) to define analysis regions at different scales.

### Overview of Models

We modeled the joint multivariate distribution of the eight leaf traits (the response variables) using three different approaches: (1) environment-only model using climate and topography as fixed effects; (2) species-only model using species as random effects, with covariances among the random effects structured by the phylogeny for all woody species recorded in the FIA database for the eastern USA; and (3) ‘combined model’ including both environmental and species effects (throughout this paper, ‘species effects’ in our models refers to phylogenetically structured species random effects). Here, we briefly summarize the modeling framework; full details are in Appendix S2. All approaches used the same joint-multilevel Bayesian framework and were evaluated using 5-fold cross-validation. The joint structure of the model allows for modeling all traits simultaneously (Wilkinson et al., 2021), considering the correlation structure across traits (i.e., inter-trait relationships, such as those typical of the LES). Environmental effects were fitted using generalized additive models (GAMs) to account for non-linear relationships. We used thin plate regression splines to estimate the smooth terms using the brms R package (Bürkner, 2017). Phylogenetic relationships across species were modeled by including species as a random effect and accounting for their phylogenetic relationships by estimating their correlation structure from cross-species cophenetic distances (Paradis et al., 2019). The distance matrix was used to estimate the correlation structure across taxa, allowing parameter estimates for the lead traits of rare or unsampled taxa to borrow strength from widely sampled species (de Villemeruil & Nakagawa, 2014). We used multivariate normal families and weakly informative priors in all cases (Appendix S2).

To reduce collinearity and the number of climate predictors, we performed a PCA for each climate variable (net radiation, precipitation, vapor pressure, maximum and minimum temperature) using monthly averages from 1985 to 2015. We used the first component of each PCA to represent each climate variable in the environment-only and combined models. To quantify uncertainty in model accuracy, we used the 95% prediction interval of the Bayesian R^2^ (Gelman et al., 2018). See additional methods details in Appendix S1-S3.

### Partitioning trait variation

We used two factors variance partitioning (Ribas et al., 2006) to estimate the relative contribution of inter-vs. intraspecific trait variation at the continental scale by comparing each single-driver model (environment-only and species-only) to the combined model following Muñoz & Real (2006). Previous studies partitioned inter-vs. intraspecific trait variation using measured trait values (Leps et al., 2011 de Bello et al., 2011). To adapt this framework to the modeling context, we quantified the proportion of variance explained (Bayesian R^2^) for each trait by each of our three models (environment, species, and combined models), and we then quantified the proportion of variance attributable to species effects alone (σ^2^_species_ = R^2^_combined_ – R^2^_env_), environmental effects alone (σ^2^_env_ = R^2^_combined_ – R^2^_species_), and joint species-environment effects (σ_joint_ ^2^ = R_combined_^2^ – σ^2^_species_ – σ^2^_env_). Note that the pure environment effect (σ^2^_env_) is a component of intraspecific variation, because σ^2^_env_ is, by definition, independent of species-level (interspecific) effects (σ^2^_species_). However, not all intraspecific variation can be attributed to measured environmental variables; thus, intraspecific variation contributes not only to σ^2^_env_, but also to unexplained variation. In summary, trait variation can be conceptualized as including pure species effects (σ^2^_species_), pure environmental effects (σ^2^_env_, which is equivalent to intraspecific variation explained by the measured environmental variables), joint species-environment effects (σ^2^_joint_), and unexplained variation (which includes measurement error as well as intraspecific variation associated with errors or omissions in the environmental data).

### Analysis methods for quantifying inter- and intraspecific variation at scale

To study inter- and intraspecific patterns in trait variation, including inter-trait relationships, we used the combined model to estimate leaf traits for all individual trees in FIA plots in the eastern USA (all USA states bordering or to the east of the Mississippi River). For each individual tree (∼1.2 million trees in ∼30,000 inventory plots), we estimated its posterior median trait values using each of our three models. For the environment-only and combined models, we used the climate principal components (see above) at the publicly reported location of each FIA plot, and we used the topographic variables (elevation, slope, and aspect) reported by FIA for each plot. Predictions from our species-only model produced broad-scale geographic patterns like those from previous species-only models (Swenson & Weiser, 2010; Clark, 2016) (Fig. S.2). We compared predictions from our species-only model (which ignores intraspecific variation) to predictions from the combined model (which includes both inter- and intraspecific variation) to explore the consequences of intraspecific variation for broad-scale distributions.

To quantify relationships between N% and LMA, we used ordinary least squares (OLS) regression slopes based on log-transformed predictions of N% and LMA from our combined model for individual trees in FIA plots. To quantify differences in N% vs. LMA relationships within and among species and plant functional types, we compared the OLS slopes from regressions based on individuals vs. species-level means, for all eastern USA species and separately for broadleaf and needleleaf plant functional types. We used OLS regression instead of standard major axis (SMA) regression because (1) this facilitated comparisons with the OLS slopes reported by Osnas et al., (2018), including inter- and intraspecific trait relationships for different species groups; (2) SMA slopes are unstable when OLS slopes are shallow, as observed for a large fraction species; and (3) we expect only small errors in the x-variable (LMA) in these regressions (see additional details on the N% vs. LMA regressions in Supplement 6).

## Results and Discussion

### Model evaluation: explained variation at continental scale

We fit the species-only, environment-only, and combined (species and environment) models using leaf trait data from NEON and evaluated their explanatory power using the Bayesian R^2^ of the predicted values for 88 out-of-sample test trees. For all 8 leaf traits, the combined model explained the largest amount of variance in the out-of-sample test data (average R^2^ across the 8 traits = 0.64), substantially outperforming both the environment-only (average R^2^ = 0.35) and species-only (average R^2^ = 0.52) models (Fig. 1, Fig. S.3). The combined model had the highest performance for predicting LMA (R^2^ = 0.81) and the lowest performance for predicting ChlBA (R^2^ = 0.51). Uncertainty in predictions was accurately estimated across all traits and for all models (Fig. S.4), with the combined model’s mean 95% coverage values ranging from 94.3% to 98.8%. Residuals from the combined model – which represent variation that is unexplained by environmental factors, species identity, or phylogeny – revealed patterns like those previously described across species, suggesting the presence of inherent trait trade-offs and correlations that occur both across species, and also within species in a given environment.

**Fig. 1.**
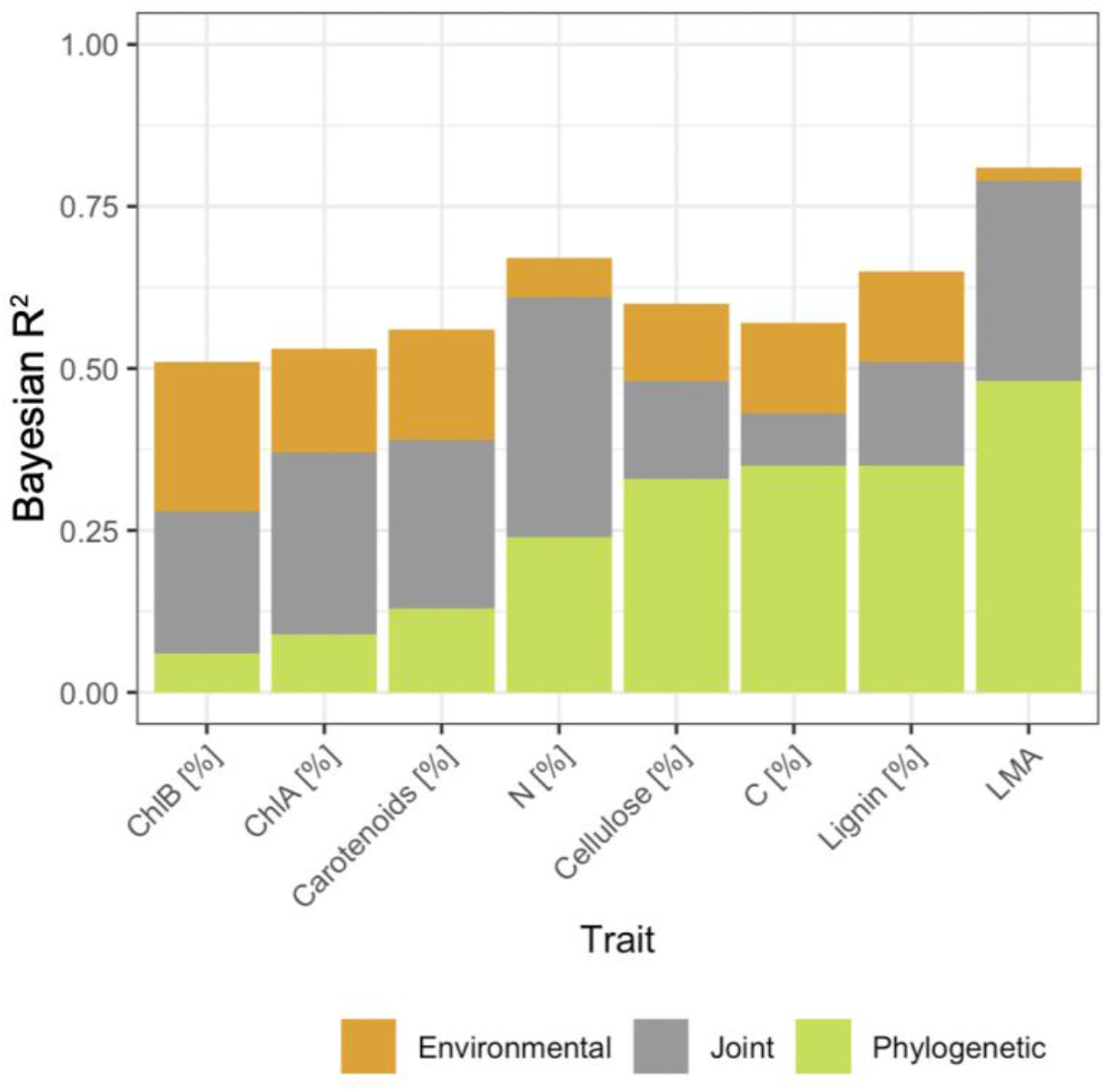
Variance partitioning between pure species effects, pure environmental effects (intraspecific variation), and joint species-environment effects based on the R^2^ for independent validation data. Pure environmental variation is independent of species variation, and vice versa. Joint variation is jointly explained by species and environmental effects and cannot be uniquely ascribed to either. LMA is leaf dry mass per area, and all other traits are % mass.

Combined-model residuals revealed a strong negative correlation between N% and LMA, as in the LES (Wright et al., 2004), and the presence of two main trait clusters (Fig. S.5): traits mainly involved in photosynthesis (N%, Carotenoids, ChlA and ChlB) and traits related to leaf structure (LMA, C%, Cellulose%, Lignin%).

### Model evaluation: geographic transferability

To determine the appropriateness of using the model to make predictions outside of the scope of the NEON dataset, we tested the performance of the combined model on novel locations and novel species using independent data from the Botanical Information and Ecology Network (BIEN) and TRY. These datasets include trait data on LMA, N% and C% for 62 eastern USA tree species, including 27 species unavailable in training data (Fig. S.1, supplement S1). The combined model showed good transferability to other data sources (mean R^2^ = 0.54, 95% coverage = 91%, Fig. S.6). Accounting for phylogenetic relationships, in addition to environmental predictors, yielded substantial model improvements when predicting traits for species not sampled at NEON sites (Fig. 2). This improvement occurs because accounting for phylogenetic structure when modeling species random effects (see Methods) allows parameter estimates for unsampled species to borrow strength from closely related species for which observations are available (Evans et al., 2016). The above transferability analyses suggest that the combined model is suitable for broad-scale applications.

**Fig. 2.**
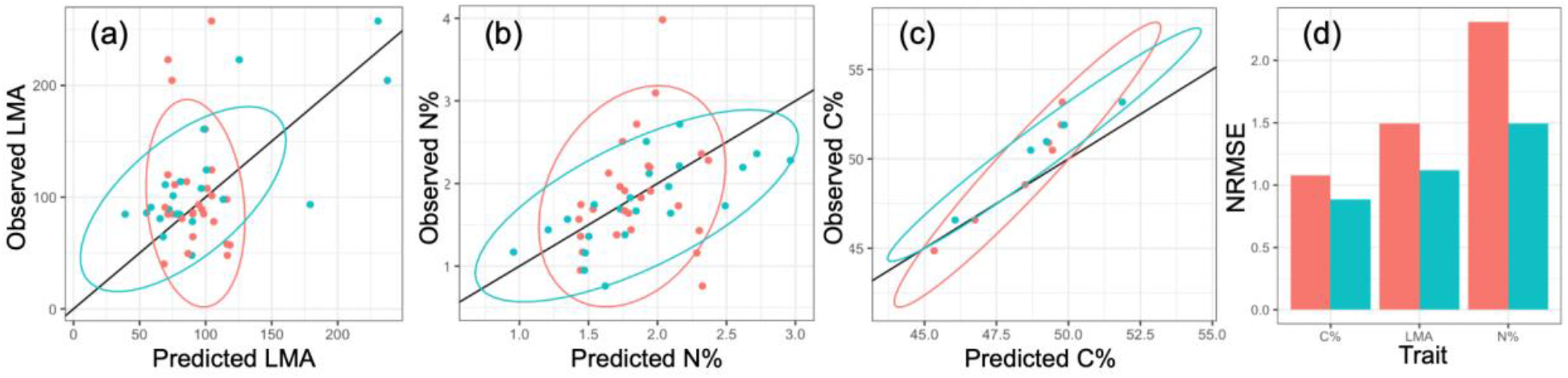
Observed vs. predicted values from the combined model for (a) LMA (g m-2), (b) N%, and (c) C%, either assuming phylogenetically-structured covariance among species random effects (azure), or assuming independent species random effects (orange). Ellipses are 95% confidence ellipses. (d) RMSE (normalized by standard deviation) showing the difference between accounting for (azure) or ignoring (orange) phylogenetic relationships when predicting traits for species not included in the training data.

### Model evaluation: intraspecific variation across wide geographic ranges

The combined model produced realistic ranges of intraspecific variation when compared with available independent data for widely distributed species (Fig. 3). Specifically, intraspecific ranges of N% predicted from the combined model were similar to those observed in field samples for three widespread and abundantly sampled species in the NEON, FIA and TRY datasets (*Abies balsamea, Acer rubrum*, and *Fagus grandifolia*). For these species (Fig. 3b-d), combined-model predictions of N% spanned an average range of 1.12 N%, which is as wide as the average difference between evergreen needleleaf and deciduous broadleaf species in the eastern USA (1.13 N% based on NEON field data). This large magnitude of intraspecific variation supports the idea that adaptation to the environment accounts for a large proportion of community-level variation and plays an important role in community-mean trait shifts across environmental gradients (Albert et al., 2010, Violle et al., 2014, Siefert et al., 2015).

**Fig. 3.**
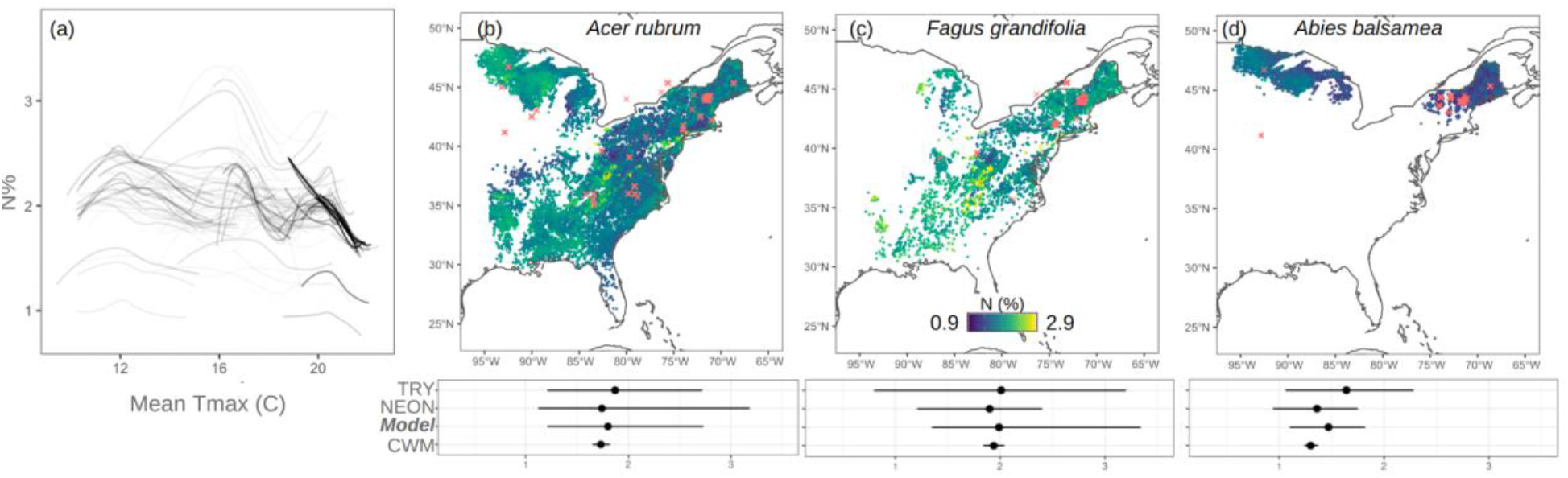
Intraspecific variation of predicted leaf N% for tree species in the eastern USA. (a) Relationships (estimated using GAMs) of N% (median predicted values) vs. maximum monthly temperature for the 91 species (each species has a separate line) showing significant effects of temperature on N% (the darkness of the curve increases with the strength of the relationship between N% and temperature). (b-d) Geographic patterns and range of intraspecific N% for two widespread deciduous broadleaf (*Acer rubrum* and *Fagus grandifolia*) and one widespread needleleaf evergreen (*Abies balsamea*) tree species. Mean (points) and 99% intraspecific ranges (lines) in b-d are from: TRY data (within the eastern USA), NEON data (within the eastern USA), the combined model, and random error from the species-only model. Red crosses represent TRY sample locations.

### Application 1: Role of species and environment in predicting traits at the continental scale

The importance of different environmental drivers varied among traits, supporting an important role of climate in driving leaf economics in local communities (Ordoñez et al., 2009). Precipitation and temperature were mainly important for traits involved in photosynthesis (N%, ChlA, ChlB and Carotenoids), generally having a positive effect on their concentration (except for ChlB, Fig. S.7). Net radiation showed a negative effect on N%, while vapor pressure generally had a negative effect on pigments but a positive effect on traits associated with leaf toughness and durability (cellulose, lignin and C%). Elevation was the most important topographic predictor and the strongest environmental driver of LMA, consistent with previous studies (Reich & Oleksyn, 2004, Poorter et al., 2009, Kitajima et al., 2016).

On average, interspecific variation (phylogenetically structured species-level effects, controlling for environmental variation; see σ^2^_species_ in Methods) accounted for 25% of the total explained variation across the 8 traits, intraspecific variation (pure environmental effects, controlling for species effects; see σ^2^_env_ in Methods) accounted for 13%, and joint species-environment effects accounted for 23% (Fig. 1). However, variance partitioning differed for photosynthetic traits (pigments and N%) vs. structural traits (cellulose%, C%, lignin%, and LMA). For photosynthetic traits, the largest variance component was the joint species-environment component (Fig. 1); i.e., for these traits, much of the explained variation was associated with coordinated shifts in the environment and species composition. In contrast, for structural traits, pure species effects (i.e., interspecific variation that is independent of environmental variation) accounted for most of the trait variation (Fig. 1). It is important to note, however, that data limitations likely lead us to underestimate the importance of pure environmental (intraspecific variation) effects. Firstly, our analysis is restricted to the upper (sun-lit) canopy. Although this is a common practice (Pérez-Harguindeguy et al., 2013), it ignores trait variation across the canopy light gradient, a major source of intraspecific variation (Lusk et al., 2008; Osnas et al., 2018). Secondly, 62% of species in our analysis were only sampled within a single NEON site. Thirdly, as in any observational study, we have only limited environmental data, which inevitably leads to underestimate environmentally structured intraspecific variation.

Despite the likely underestimation of intraspecific variation in our analysis, pure environmental effects (a component of intraspecific variation; see Methods) accounted for a substantial fraction of explained variation for all traits except for N% and LMA (Fig. 1). Thus, although species-only models accounted for the majority of explained variation (i.e., the sum of the joint effects and species effects in Fig. 1) – consistent with previous studies highlighting the important role of species distributions in predicting traits at broad geographic scales (Siefert, et al., 2015; Clark 2016, Yang & Swenson, 2018) – predictions for most traits can be substantially improved by a combined species-environment modeling approach. This conclusion – i.e., that accounting for intraspecific variation can enhance trait predictions − would likely be strengthened by improved data on traits and environmental variables.

### Application 2: Drivers of ecoregional differences in traits distribution

Predictions from the combined model show broad-scale patterns associated with shifts in forest communities (Fig. S.8) and large-scale climatic and topographic patterns across latitudinal and altitudinal gradients (Fig. 4). In some cases, trait distributions shifted abruptly between neighboring ecoregions due to a combination of shifts in local environmental conditions and in community assembly (independent of the environment). For example, changes in species composition may explain trait differences between the Mississippi Alluvial Plains and neighboring ecoregions (Fig. S.9). The Mississippi Alluvial Plains ecoregion is characterized by agricultural land use, and forests are often limited to riparian ecosystems favoring bottomland broadleaf species (e.g., *Celtis laevigata, Fraxinus pennsylvanica, Salix nigra*) in contrast to needleleaf species (e.g., *Juniperus virginiana, Pinus taeda*, and *Pinus echinata*) more common in the neighboring ecoregions. Being that broadleaf species are generally characterized by higher N% and lower LMA, this change in community assembly translates into predicted regional trait patterns. Other ecoregion boundaries show little change in species composition, in which case geographic variation in traits may be attributed to environmental effects on intraspecific variation. For example, in the Appalachian Mountains, opposing patterns are exhibited at high altitudes in the Blue Ridge (lower N% and pigment; higher LMA and C%) compared to valleys piedmont in the neighboring ecoregions (higher N% & pigments, lower LMA & C%) (Fig. S.10).

**Fig. 4.**
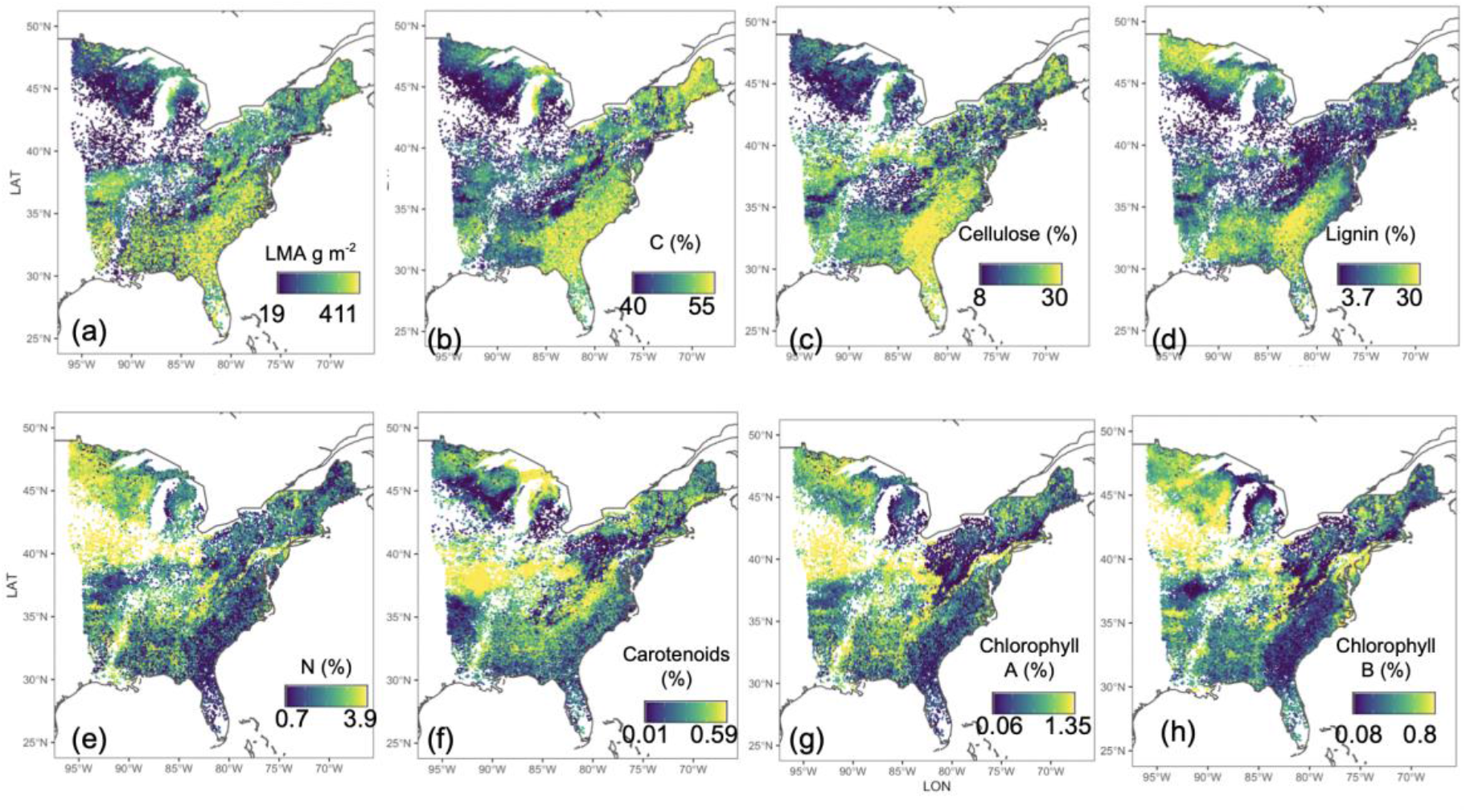
Distributions across the eastern USA for leaf structural (LMA, lignin, cellulose and C%), macro-nutrient (N%), and pigment (carotenoids, chlorophyll A and B) traits. Values represent trait averages at each FIA plot (n = 30,331 plots), calculated across all individual trees sampled within the plot.

Predictions from the combined model differed from the species-only and environment-only models, with the direction and magnitude of divergences varying across ecoregions (Fig. 5, Fig. S.11). Species-only and environment-only model predictions differed significantly from the combined model (p < 0.0001 in paired t-tests) in 80% cases (different combinations of traits, ecoregions, and models). Similarly, predictions from the species-only and environment-only models differed from each other (p < 0.0001 in paired t-tests) for 93% of ecoregion-trait combinations, which demonstrates the distinct effects of species and environmental drivers on trait distributions and shows the importance of a combined approach for prediction & inference.

**Fig. 5.**
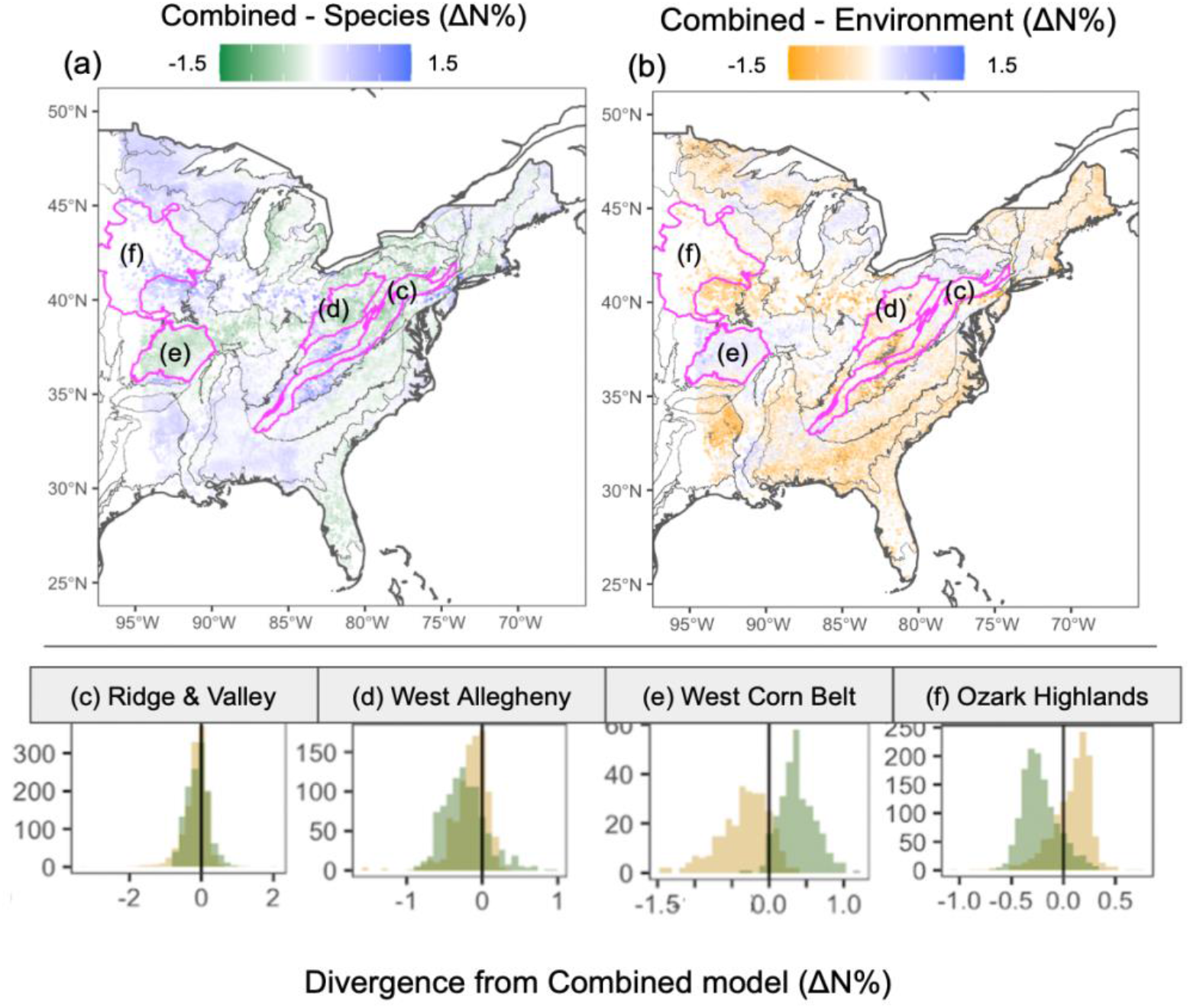
Divergence in N% between the combined model and the species-only (a) and environment-only (b) models. Dots represent FIA plots where predicted N% was higher in the combined (blue), environment (orange), or species (green) model. White areas represent plots with little to no divergence (p > 0.0001), or areas with no FIA plots recorded. (c-f) Examples of divergence patterns in four ecoregions (see the outlined ecoregions in the maps, labeled c-f according to their corresponding histogram): combined model minus environment model (orange) and combined model minus species model (green). Vertical black lines represent zero divergence. Ecoregion examples include: all three models capturing similar information with ΔN% near 0 (c); species and environment models similar to each other but diverging from the combined model (d); and and divergence among all three models (e-f).

Patterns of divergence from the combined model indicate how species and environmental effects vary geographically across the eastern USA (Fig. 4). In regions with no significant divergence (e.g., the Ridge and Valley region), species-only and environment-only models yield similar results to each other and to the combined model. This convergence suggests a limited role for intraspecific variation combined with strong environmental filtering of species assembly. However, for most ecoregions, species-only and environment-only models diverge from the combined model in opposite directions (i.e., positive divergence for one and negative divergence for the other) (Fig. 5d, 5e, 5f, S.11). Significant divergence for the species-only model indicates that continental-scale species averages fail to correctly represent trait values at finer scales, due to local environmental conditions and/or competition shifting their values away from the mean.

Areas where the combined model predicts higher N% than the species-only model (blue shading in Fig. 5a) suggest the presence of environmental effects that increase N% above species means. Conversely, orange-shaded areas in Fig. 5a indicate environmental effects that lower N% below species means. These patterns may be explained by regulatory mechanisms adjusting allocation to proteins, pigments, or structural compounds to balance photosynthetic capacity, toughness, and chemical defense across environmental gradients (Weih & Karlsson, 2001, Tjoelker et al., 2001, Albert et al., 2010). In contrast, divergence between the combined and environment-only models (Fig. 5b) could be explained by (i) environmental effects that have a phylogenetic signal not captured by the environmental variables included in our analysis; and/or (ii) stochastic factors (e.g., disturbance history and dispersal limitation) that have resulted in species distribution patterns that are decoupled from current environmental conditions (Burns & Strauss, 2012, McIntyre et al., 1999). Testing mechanistic hypotheses for the above divergence patterns is beyond the scope of our study. Nevertheless, identifying these biogeographic patterns is a step towards better understanding trait distributions and is only possible using modeling approaches that combine phylogenetic and environmental information.

### Application 3: Large scale intraspecific trait variation driven by environmental drivers

In addition to mapping species and environmental trait signals, another essential reason for a combined approach is that it allows for predictions of intraspecific variation for individual species (Fig. 3). Models based on community-weighted means, by definition, do not account for intraspecific variation. In contrast, environment-only models implicitly capture at least some component of intraspecific variation (see Methods) but cannot separate this intraspecific variation from trait variation due to shifts in species composition (Hulshof & Swenson, 2010). Our combined modeling approach makes this separation possible for hundreds of species-trait combinations. Quantifying intraspecific variation in a comprehensive manner (i.e., for all species across a large geographic region) is a necessary step towards testing hypotheses about the role of individual trait variation in species coexistence and interspecific competition (Hart et al., 2016). Here, we explore the potential for using a combined species-environment approach to studying intraspecific variation by evaluating how its distribution vary among eastern USA species.

Predicted patterns of intraspecific variation varied widely across species. For N%, the ratio of predicted intraspecific to total observed interspecific variation ranged from less than 10% for species with limited geographic ranges (e.g., *Populus heterophylla, Sabal palmetto*, and *Gleditsia aquatica*) to over 60% for broadleaf species with broad geographic ranges (e.g., *Cercis canadensis, Betula lenta*, and *Carpinus caroliniana*). Intraspecific responses of N% to temperature also showed a variety of patterns among species (Fig. 3a, Fig. S.12), including (1) bell-shaped patterns (n = 85) like those observed in compilations of field data (Reich & Oleksyn, 2004, Laughlin et al., 2012); (2) negative relationships (n = 7); and (3) cases with no significant relationship (n = 108). Note that our modeling approach does not explicitly include species-by-environment interactions. Thus, differences among species in their magnitudes of intraspecific variation and in their trait-environment relationships emerge from how species distributions in environmental space combine with non-linear trait-environment relationships.

### Application 4: Inter- and intraspecific trait relationships: implications for the leaf economics spectrum and terrestrial ecosystem models

In terrestrial ecosystem models, accurately representing leaf functional diversity (e.g., biogeochemical and/or land-surface models) requires capturing variation within and among species (Bonan et al., 2002, Osnas et al., 2018). The LES provides a simple, one-dimensional framework for understanding leaf functional variation from local to global scales (Wright et al., 2004) and for representing plant functional diversity in terrestrial ecosystem models (Sakschewski et al., 2015), including the land components of Earth System Models (Bonan et al., 2002, Fisher et al., 2015). On the other hand, divergent patterns in trait variation observed within species, among species, and among functional or taxonomic groups (Osnas et al., 2018) and at different taxonomic scales (Anderegg et al., 2018) suggest that a one-dimensional framework may be too simplistic. There is growing interest in enhanced representations of trait diversity in terrestrial ecosystem models (Fisher et al., 2015, Sakschewski et al., 2015, Dantas de Paula et al., 2021), but these model developments are currently data-limited due to limited information on trait distributions at broad geographic scales (see details below).

When expressed per-unit leaf mass, leaf traits related to photosynthesis, metabolism, and nutrient concentrations typically have strong negative covariance with LMA across species (Wright et al., 2004, Lloyd et al., 2013, Osnas et al., 2013) but are often roughly invariant within species across canopy light gradients (Ellsworth and Reich 1993, Evans and Poorter 2001, Niinemets et al., 2015). Constant mass-normalized traits across canopy light gradients − equivalent to a roughly linear increase in the area-normalized value of these same traits as LMA increases from shade to sun (Osnas et al., 2018) − is accounted for in some terrestrial ecosystem models (e.g., Sellers et al., 1992, Bonan et al., 2012). Intraspecific variation in leaf traits across broad environmental gradients can be substantial as well, with important consequences for ecosystem models (Reich et al., 2014, Dong et al., 2017). Our broad-scale systematic mapping of N% and LMA, for individual trees (summarized at the FIA plot level in Fig. 4, Fig. S.13) enables further understanding of intraspecific trait relationships across geographic space.

For the eastern USA, our results show that slopes of N% vs LMA within and among species fall between the shallow slopes found within species across light gradients (typically between 0 and −0.3; Supplement 6), and the steeper slopes recorded among global species (−0.6; Supplement 6). These results were consistent for both deciduous broadleaf (DB) and evergreen needleleaf (EN) plant functional types (PFTs; Table S.2). Specifically, in our analysis, intraspecific slopes had median values around −0.37 (Fig. 6a-b), slopes across all individuals within PFTs were −0.36 for DB and −0.48 for EN (Fig. 6c), and slopes across species means were −0.4 for DB and −0.45 for EN (Fig. 6d). These intermediate values (i.e., steeper than those observed across intraspecific light gradients, but shallower than observed across global species) may be explained by partially compensating factors. LMA variation associated with intraspecific LL variation across space may largely involve variation in low-nutrient, structural components of LMA (as occurs across species; Kitajima et al., 2021), which would lead to steep N% vs. LMA slopes (Osnas et al., 2018). LMA variation associated with certain climatic responses (e.g., sensitivity of leaf water potential to tissue water content; Niinemets 2015) may also largely involve low-nutrient structural mass, which would again lead to steep N% vs. LMA slopes. In contrast, water loss can be reduced at dry sites (which tend to have high LMA) if N% and thus photosynthetic efficiency are also high (Wright et al., 2001), which would lead to positive N% vs. LMA slopes across moisture gradients. This combination of mechanisms, and perhaps similar mechanisms operating along geographic gradients (e.g., temperature extremes & season length), may result in intraspecific leaf trait relationships across space that are intermediate to those driven by photosynthetic responses to light vs. those driven by interspecific variation in LL.

**Fig. 6.**
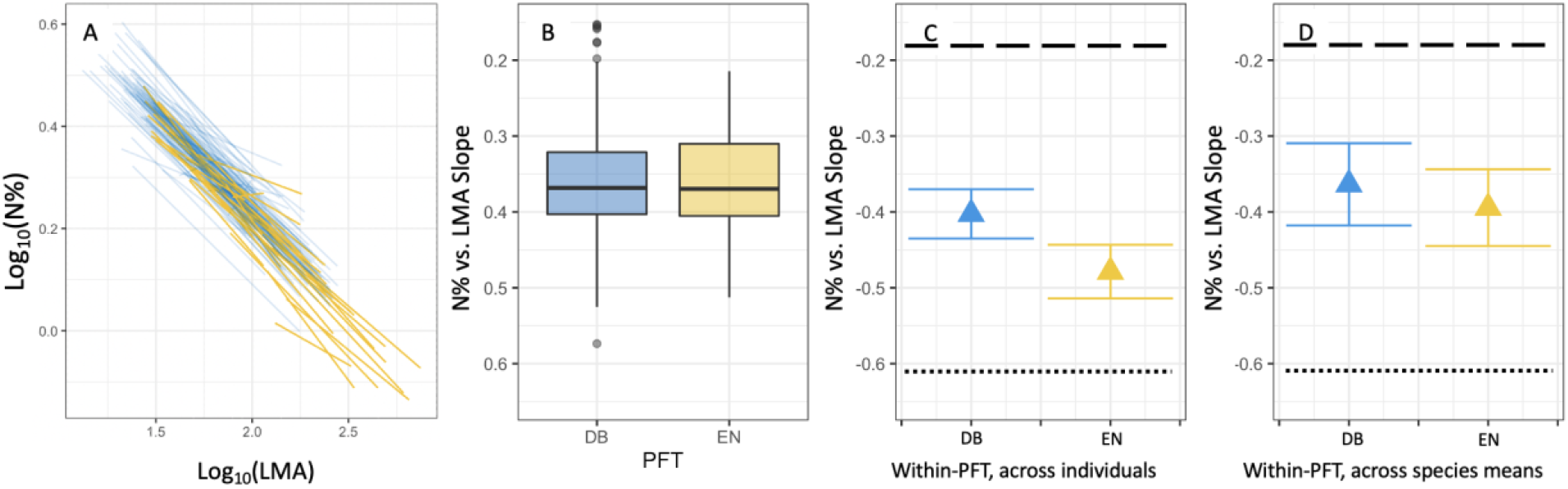
Relationships between leaf nitrogen percent mass (N%) and leaf mass per area (LMA) across the eastern USA, based on combined-model predictions of N% and LMA for individual trees in eastern USA Forest Inventory and Analysis (FIA) plots. (A) Fitted intraspecific ordinary least squares (OLS) regression lines for eastern USA deciduous broadleaf (DB: blue) and evergreen needleleaf (EN: orange) species. Each line is an individual species. (B) Distribution of intraspecific OLS slopes for DB and EN species; bars are medians, boxes span the interquartile range, whiskers span 95% of slopes, and points show outliers. (C) OLS slopes (± s.e.) across all individuals within each PFT. (D) OLS slopes (± s.e.) across species means within each PFT. Individual trait values were log_10_-transformed for analyses in (A-C), and species means were log_10_-transformed for analyses in (D). Dashed and dotted black lines in panels C and D are from Osnas et al., (2018): the dashed line at −0.18 represents the intraspecific slope across the canopy light gradient, and the dotted black line at −0.62 represents the slope across global species (see details in Supplement 6).

Our results suggest that to accurately represent leaf functional variation, terrestrial ecosystem models may need to consider three different types of variation: intraspecific variation across light gradients, intraspecific variation across geographic environmental gradients, and interspecific relationships (which may differ among PFTs). Given the impracticality of parameterizing species-level global ecosystem models, we expect these models to continue to explore approaches to representing trait variation within broadly defined PFTs without explicitly modeling individual species (e.g., Fisher et al., 2015, Sakschewski et al., 2015). Accurately representing the different intra- and interspecific patterns described above does not necessarily require an explicit species-level approach. A greater understanding of the anatomical and physiological mechanisms underlying trait variation (e.g., Niinemets et al., 2015) could lead to a generalized, species-independent model that captures divergent patterns within & among PFTs.

Our results have implications for interpreting interspecific leaf trait relationships. PFT-level slopes across individuals (Fig. 6c) differed from slopes across species means (Fig. 6d). These differences in slope are greatly reduced when species are rarified to equal abundance (Fig. S.14), suggesting that differences in slope are largely due to differences in species abundance: more common species exert greater influence in the individual-level analysis (Fig. 6c), whereas all species contribute equally to the species-level analysis (Fig. 6d). Therefore, accounting for within-species relationships may be relevant for some specific applications. For example, when parametrizing or testing ecosystem models, trait relationships estimated from individual-level datasets (that preserve species differences in abundance and intraspecific variation) should more accurately capture ecosystem-level responses compared to analyses based on species means.

## Conclusions

Both phylogenetic and environmental effects are fundamental to understanding drivers and distribution of plant traits. Combining both in a single model is challenging due to data limitations but is possible by leveraging large scale datasets. This approach allows for improved traits predictions compared to models that rely on either species means or environmental conditions in isolation and allows for robust predictions for species and regions not included in the training data. Across the eastern USA, interspecific trait variation, geographic variation in species’ abundances, and intraspecific variation all have important effects on trait distributions and LES relationships. The influence of these drivers varies by species, trait, ecoregion, & scale.

Our approach overcomes data limitations by integrating multiple sources of biological and environmental information to create a single integrated model. As new traits, phylogenetic, and vegetation inventory data become available globally, combined modeling approaches, such as the one presented here, can be applied to other geographic regions to study intraspecific variation and trait-species-environment relationships. Such analyses are already possible using national forest inventories in some regions (e.g., Canada, New Zealand & European countries; Schelhaas et al, 2006, Gills et al., 2005, Paul et al., 2021) in combination with ever-growing plant trait data (e.g., TRY; https://www.try-db.org/). Expanding this work beyond the USA may contribute to further understanding the mechanisms driving trait distributions across scales and the link between traits, species distributions, community assembly, and ecosystem function.

## Acknowledgements

This work was supported by the Gordon and Betty Moore Foundation’s Data-Driven Discovery Initiative grant GBMF4563 to E. P. White; National Science Foundation (NSF) grant 1926542 to E. P. White, S. A. Bohlman, A. Zare, D. Z. Wang, and A. Singh; NSF grant DEB-1442280 to J. W. Lichstein and S. A. Bohlman; USDA/NIFA McIntire-Stennis program grant FLA-FOR-005470 to S. A. Bohlman; University of Florida Biodiversity Institute (UFBI) and Informatics Institute (UFII). No additional funding was received for this study.

## Statement of authorship

SM, JL and EW designed the experiment and the methods; SM performed the analysis; EW, AS, SB and BW supervised the work, helped with experimental design, and supervised analysis and technical aspects. All authors contributed to the manuscript.

