## Supplementary materials for "Disentangling the roles of inter- and intraspecific variation on leaf trait distributions across the eastern United States"

**Supplement1: Data preprocessing**

**S1.1 NEON data**

Trait data from the National Ecological Observatory Network (NEON) on nitrogen (N%), carbon (C%), chlorophyll A (ChlA%), chlorophyll B (ChlB%), carotenoids (Crt%), leaf mass per area (LMA, gm^-2^), lignin (%) and cellulose (%)) were downloaded from the neonUtilities R package (Lunch et al., 2020). Raw trait data were aggregated using the functions in neonVegWrangleR (https://zenodo.org/badge/latestdoi/215943021). Since carotenoids and chlorophyll were not normalized per mass %, we converted their values using the following formulation:

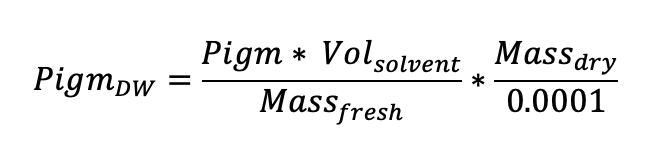

Only trait data collected after 2014 was used due to its higher quality. We did not use trait data from the Mountain Lake Biological Station (MLBS) NEON site due to issues with laboratory measures for carbon and microelements (NEON staff personal communication). We did not include micronutrients in the model due to low sample sizes after these restrictions to the dataset. When multiple leaf samples from a single individual tree were available they were averaged to a single vector of traits for that individual. We only included individuals for which data on all 8 traits were available. GPS coordinates of individual trees were calculated using the approach in https://github.com/weecology/neonVegWrangleR. Terrain data (elevation, slope and aspect) were extracted using NEON AOP data. Aspect and slope were transformed using a sin-cos transformation to rescale their variation into the same range as the other environmental features.

**S1.2. Transferability dataset**

Transferability analysis was performed using data from Botanical Information and Ecology Network (BIEN) and a subset of the TRY dataset (Enquist et al., 2009, Kattge et al., 2020). For BIEN, we extracted all data from the region of interest, between 27 and 45 degree latitude, and -95 and -55 degrees longitude. We focused only on traits from species of shrubs or trees. Data was downloaded from the R package BIEN (Maitner, 2020).

Data from TRY consisted of individual traits data for C%, N% and LMA from the OSBS, TALL and MLBS NEON sites (Marconi et al. 2021). Separate NEON data from only one of these sites (TALL) was used in the model development. Data from this dataset was collected to maximize the observed range of traits, and was fairly balanced between common and rare species. From this dataset, we used only sunlit leaves. For both datasets, topographic features (elevation, slope and aspect) were extracted from USGS 3DEP viewer (<https://apps.nationalmap.gov/3depdem/>).

Unknown species were predicted by making predictions for the 3 closest species in the phylogenetic tree and then taking the average, weighted by the cophenetic distance of these three species from the targeted species.

Another subset of TRY was used to compare the range of N% variability of *Acer rubrum* (n = 80), *Fagus grandifolia* (n = 76), and *Abies balsamea* (n = 130) observed from the field with predicted intraspecific variability. For this analysis, we used only N% data from the open access portion of the TRY 5.0 database (i.e. data not subject to embargo, as defined by TRY commonly agreed [Intellectual Property Guidelines](https://www.try-db.org/TryWeb/TRY_Intellectual_Property_Guidelines.pdf), <https://www.try-db.org/TryWeb/TRY_Intellectual_Property_Guidelines.pdf>). We removed all data recorded as being from for senescent leaves, for which there were no geographic coordinates, or that could not be normalized to mass percent.

**S1.3. FIA data**

We used the most recent data on species abundances for each FIA plot, which includes survey campaigns occurring from 2016 to 2019. We used the jittered latitude and longitude data that are publicly available for use. Since terrain information is more prone to noise if extracted from the jittered coordinates, we used terrain data recorded directly in the FIA dataset. Elevation was converted to meters. Aspect and slope were transformed using a sin-cos transformation to rescale their variation into the same range of the other environmental features.

**S1.4. Phylogenetic matrix**

Phylogenetic data was collected by querying the oOpen tTree of Llife (OTL, Redelings et al., 2017) from the rotl package (Michonneau et al., 2016). We extracted the phylogenetic tree for all the species recorded in the FIA dataset. For those species absent in the OTL, we used their genus (*Rubus*, *Ceanothus*, *Crataegus*, *Prunus* , *Colubrina*). To create the phylogenetic correlation matrix, we calculated the cophenetic distance (pairwise distances between the pairs of tips from the phylogenetic tree) using the ape package (Paradis E. & Schliep K. 2019). Since the correlation matrix was not positive definite, we computed the nearest positive definite matrix to approximate the final phylogenetic correlation matrix.

**S1.5 Climate data**

Climate variables from Daymet (Thornton et al., 2018) were downloaded using the daymetR (Hufkens et al., 2018) package using the coordinates of each individual tree. We downloaded a 30 year daily time series of minimum and maximum temperature, vapour pressure, net radiation and precipitation. We discarded day length due to its direct relationship with latitude to avoid possible artifacts in the analysis of intraspecific variation of traits across a latitudinal gradient. We discarded snow-melt because of a large number of artifacts in this measure observed at FIA plot locations. Daily values were averaged to monthly time series, and further aggregated into one single vector per climate feature, thus producing a climate data table of one value of minimum and maximum temperature, vapour pressure, net radiation and precipitation for each individual tree.

**Supplement 2: Climate PCA transformation**

Daily time series of climate variables were transformed into a single value per individual tree per climatic feature (i.e. maximum temperature, minimum temperature, vapour pressure, precipitation and net radiation). Daily values were first aggregated to monthly averages over 30 years (1985 to 2015). For each climatic feature, we performed a PCA transformation and extracted the first principal component (Benestad et al., 2015). We used data from FIA plot locations to build the PCA, because (1) FIA plots likely hold most of the range of variation for each climatic feature, and (2) it represented the most general solution for scaling and centering in zero (prerequisite of PCA). To transform climate time series for NEON, TRY and BIEN data, we first centered them using the scaling factors calculated from the FIA dataset, and applied each climate feature’s specific PCA transformation derived from the FIA plot locations. PCA regression and transformations were performed using the caret package (Kuhn, 2020)

**Supplement 3: The hierarchical joint Bayesian model structure**

All models developed in this project belong to the family of multilevel bBayesian joint models (also known as hierarchical bBayesian joint models). This class of models offers the flexibility to leverage the relationships among data grouped into one or more nested categories (e.g. species classes). The core of these models is the prediction of a vector of responses (the traits) through the linear combination of environmental predictors (population effects) and varying species specific parameters (grouping effects) (Box S1). Fixed effects (parameters at the level of population) were standardized with mean of in 0 and by their standard deviation. We chose weakly informative priors for these features, with a mean of 0 and standard deviation of 1. Grouping effect (i.e. tree taxa) followed a multivariate normal distribution with mean 0 and covariance matrix 𝛴. To take phylogenetic relationships into account, the covariance matrix was calculated as the Kronecker product between the covariance matrix Vk and the known covariance matrix of phylogeny (Ak). We estimated the phylogenetic covariance matrix by using OTL cophenetic distance. Vk was parameterized using a LKJ correlation prior and a vector of standard deviation 𝝈k. To estimate all traits jointly, we used a multivariate normal distribution family. Since the responses (traits) are not independent, we added a correlation term to take into account their relationships. This term was calculated from the correlation between the residuals of each of the modeled responses as part of the fitting process. Models were built using the brms package (Bürkner, 2017). This package uses STAN for estimating the posterior distribution (http://mc-stan.org/). Estimates of the marginal distribution of fixed effects were generated by sampling the posterior of the combined model (Bürkner, 2017).

To reduce computational demand and storage space, we (1) used models trained and evaluated on a single train-test split, (2) made predictions from 200 samples rather than the entire posterior distribution,; (3) stored only means, 95 prediction intervals and their ranges, and (4) predicted unknown species by making predictions for the 3 closest species in the phylogenetic tree and then took the average weighted by the cophenetic distance of these three species from the targeted species (rather than sampling the entire phylogenetic tree). The choice of using a single train-test split was driven by its significantly lower demand in computational resources compared to a cross-validated approach. Specifically, using a 5-fold cross validated model for making predictions at FIA would result in ~15x times more computational expense for the already very slow prediction step (due to the need to sample the posterior from each fold). To demonstrate that results from a single train-test split are still robust compared to the more favorable but computationally demanding cross validation, we performed a new 5-fold cross validation for the modeling fitting and evaluation (Table S.1).

**Supplement 4: From variance partitioning to estimates of intra-inter species variation**

Variance uniquely explained by phylogenetic effects is independent is by definition independent of environmental effects and thus represents pure interspecific variation. Similarly, variance uniquely explained by environmental drivers is independent of phylogenetic effects (i.e., shifts in species composition) and thus represents intraspecific trait variation. Variance explained by the combined phylogenetic-environment model but not uniquely ascribable to either phylogeny or environmental effects may include variation that could be explained by either phylogenetic or environmental effects alone, as well as variation that can only be explained by a combined approach. Variance explained by non-combined approaches (i.e., phylogeny-only or environment-only models) is difficult to interpret because these approaches ignore the joint variance component; thus, they do not allow for variance partitioning and may therefore overestimate the pure phylogenetic and pure environmental variance components.

**Supplement 5**

<https://docs.google.com/spreadsheets/d/1gdpUzFdYaql7ujH7VIrs5xin17CtNqTCK7TJQgDsgNc/edit#gid=291899389>

**Supplement 6: Estimating and comparing N% vs. LMA slopes**

To quantify relationships between nitrogen percent leaf mass (N%) and leaf mass per area (LMA), we used ordinary least squares (OLS) regression slopes based on log_10_(N%) and log_10_(LMA). Some analyses of leaf traits employ standard major axis (SMA) regression, which accommodates errors in both x and y variables. We opted to use OLS regression slopes for several reasons: (1) this facilitated comparisons with the OLS slopes reported by Osnas et al. (2018), including intra- and interspecific trait relationships for different species groups; (2) SMA slopes are unstable when OLS slopes are shallow (as occurs in some cases; see main text); and (3) LMA (the x variable in our regressions) is relatively straightforward to measure, so we do not expect large errors in x in our analysis.

To compare our slopes for N% to those reported by Osnas et al. (2018) for nitrogen per-unit leaf area, we calculated *b* − 1 and *w* − 1, respectively for the between-species (*b*) and within-species (*w*) slopes reported by Osnas et al. (2018), which converts their leaf-area-based slopes to leaf-mass-based slopes (see their equations 2-3 and 5-6). For the global interspecific log(N) vs. log(LMA) slope, Osnas et al. (2018) report *b* = 0.38 for N per-unit leaf area (see their Table S4), which is equivalent to *b* – 1 = −0.62 for N%. For the intraspecific log(N) vs. log(LMA) slope across canopy light gradients, Osnas et al. (2018) report *w* = 0.7 and 0.94 for N per-unit leaf area at two tropical forest sites (see their Table S3), which is equivalent to *w* – 1 = −0.3 and −0.06 for N% (or −0.18 on average), consistent with the roughly constant N per-unit leaf mass values within species across light gradients reported for a number of sites across global terrestrial biomes (Ellsworth and Reich 1993, Evans and Poorter 2001, Thornton and Zimmermann 2007, Niinemets et al. 2015). Although some studies report significant relationships between N% and LMA across intraspecific light gradients (e.g., Reich and Walters 1994), these mass-based relationships are much weaker than those based on N per-unit leaf area, which again supports the rough invariance of N% across intraspecific light gradients and contrasts with patterns observed across diverse species assemblages (strong mass-based and weak area-based relationships; Reich et al. 1997, Wright et al. 2004, 2005, Osnas et al. 2018).

| 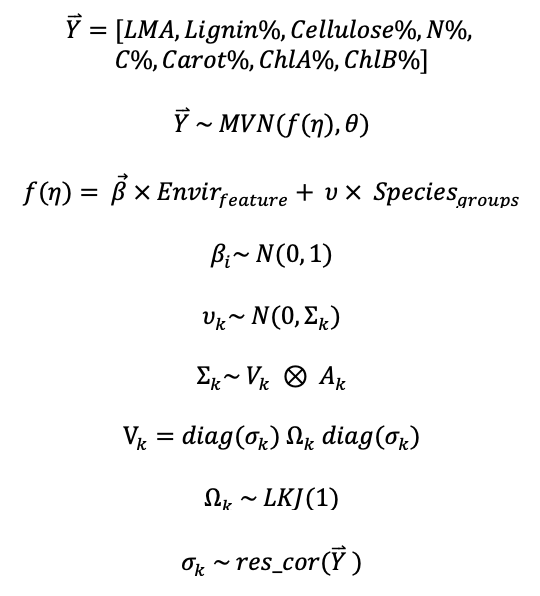 | - *θ* is the variance-covariance matrix at the population level;  - *β* are the population parameters (also known as fixed effects);  - *Envir features* are the environmental fixed effects;  - *i* is the ith feature;  - *u* are the grouping parameters (also known as random effects);  - *Species groups* are the grouping effects;  - *Σ_k_* is the grouping effect covariance matrix;  - *A_k_* is the phylogenetic covariance matrix;  - *V_k_* is the grouping effects parameters covariance matrix;  - *Ω_k_* is an LKJ correlation prior;  - *σ_k_* is the vector of responses standard deviation;  *- k* is the kth species. |
| --- | --- |

**Box S1. Core equations of the combined model.**

| Combined model | | |  | Species only | | |  | Environment Only | | |
| --- | --- | --- | --- | --- | --- | --- | --- | --- | --- | --- |
| Single fold | | 5fold-cv |  | Single fold | | 5fold-cv |  | Single fold | | 5fold-cv |
| N% | 0.77 | 0.67 |  | N% | 0.71 | 0.61 |  | N% | 0.47 | 0.43 |
| C% | 0.68 | 0.65 |  | C% | 0.54 | 0.51 |  | C% | 0.29 | 0.30 |
| lignin% | 0.61 | 0.57 |  | lignin% | 0.44 | 0.43 |  | lignin% | 0.22 | 0.22 |
| cellulose% | 0.58 | 0.60 |  | cellulose% | 0.52 | 0.48 |  | cellulose% | 0.23 | 0.27 |
| LMA | 0.84 | 0.81 |  | LMA | 0.81 | 0.79 |  | LMA | 0.34 | 0.33 |
| chlA% | 0.53 | 0.53 |  | chlA% | 0.39 | 0.37 |  | chlA% | 0.39 | 0.44 |
| chlB% | 0.56 | 0.51 |  | chlB% | 0.33 | 0.28 |  | chlB% | 0.45 | 0.45 |
| crtnd% | 0.59 | 0.56 |  | crtnd% | 0.46 | 0.39 |  | crtnd% | 0.40 | 0.43 |

Table S.1. Comparison of the three models’ evaluation resultsmodels’ evaluation between a single stratified train-test split and a 5-fold cross validation.

**Supplementary images**
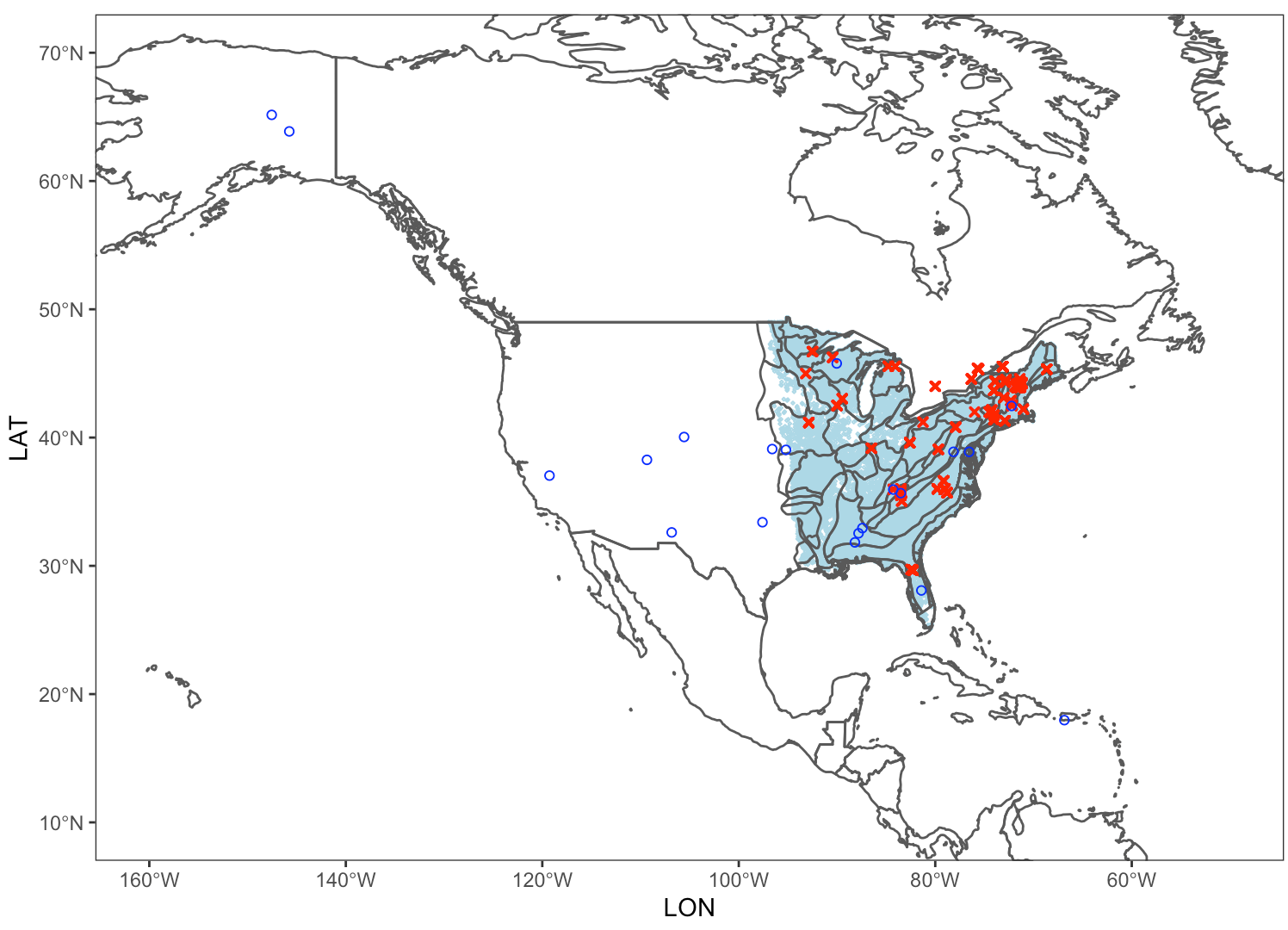

Figure S.1. Geographic distribution of data per dataset. Blue dots are the locations from NEON dataset (20 sites, n = 477 trees, train = 389 trees, test = 88 trees); cyan dots the locations of eastern USA FIA plots (n ~1.2M trees in ~30,000 plots), red x’s the locations of the external trait dataset (BIEN & TRY) used to test transferability of the combined model (n = 353 trees).

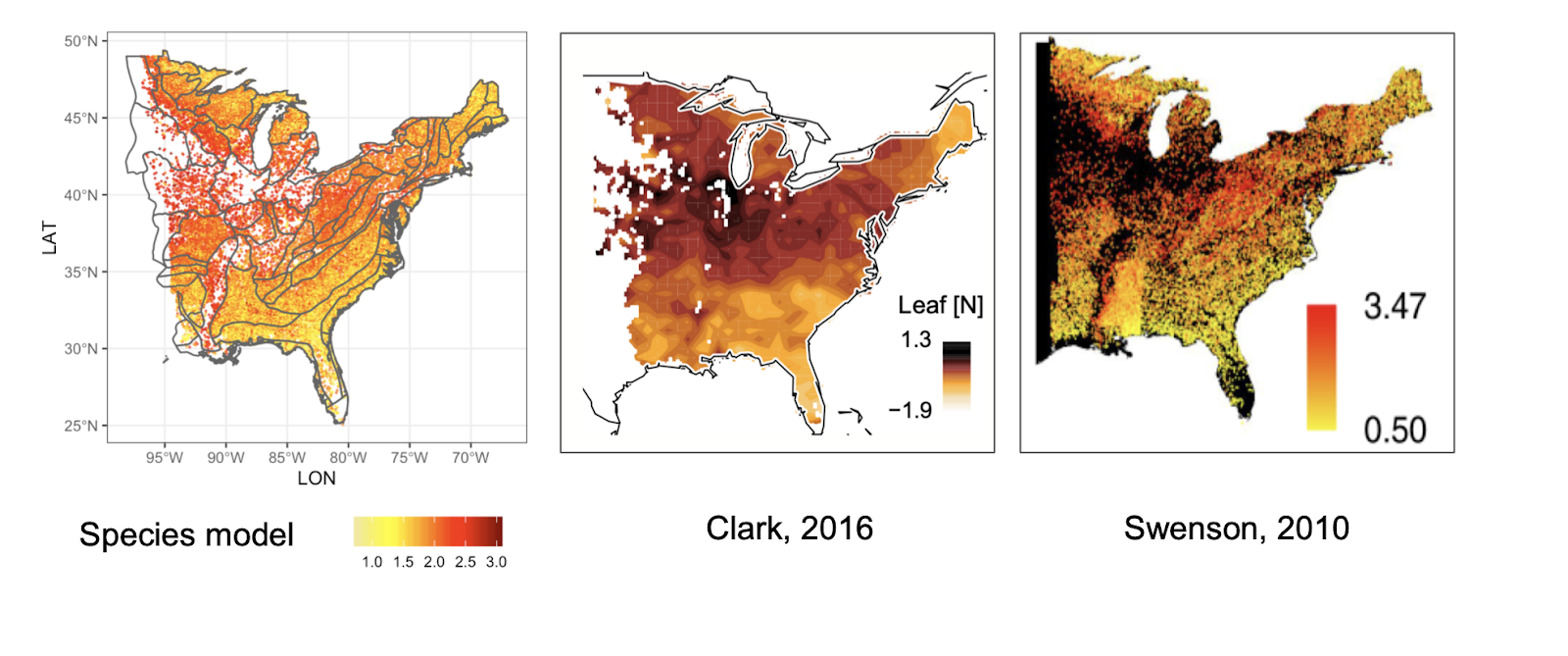

Figure S.2. Comparison between recent species distribution studies using community weighted averages from (a) the phylogeny-only model from this study, (b) Clark et al. (2016), and (c) Swenson & Weiser (2010).

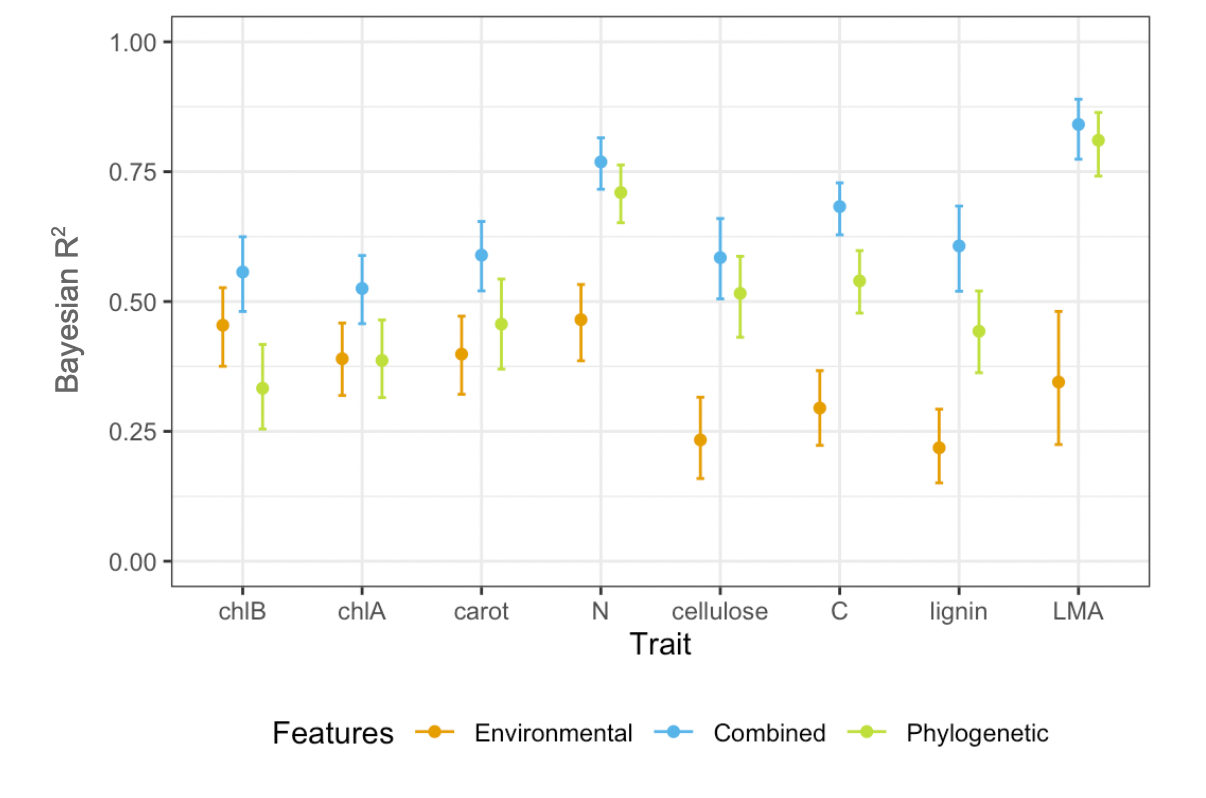

Figure S.3. R^2^ for each trait for the validation dataset (i.e., out-of-sample test data) from NEON sites (n = 88) with associated uncertainty. 95 prediction interval range of Bayesian R^2^ for each trait. Orange is the variance explained by the environment only model, green by the species only, and blue is the portion of variance explained by the combined model.

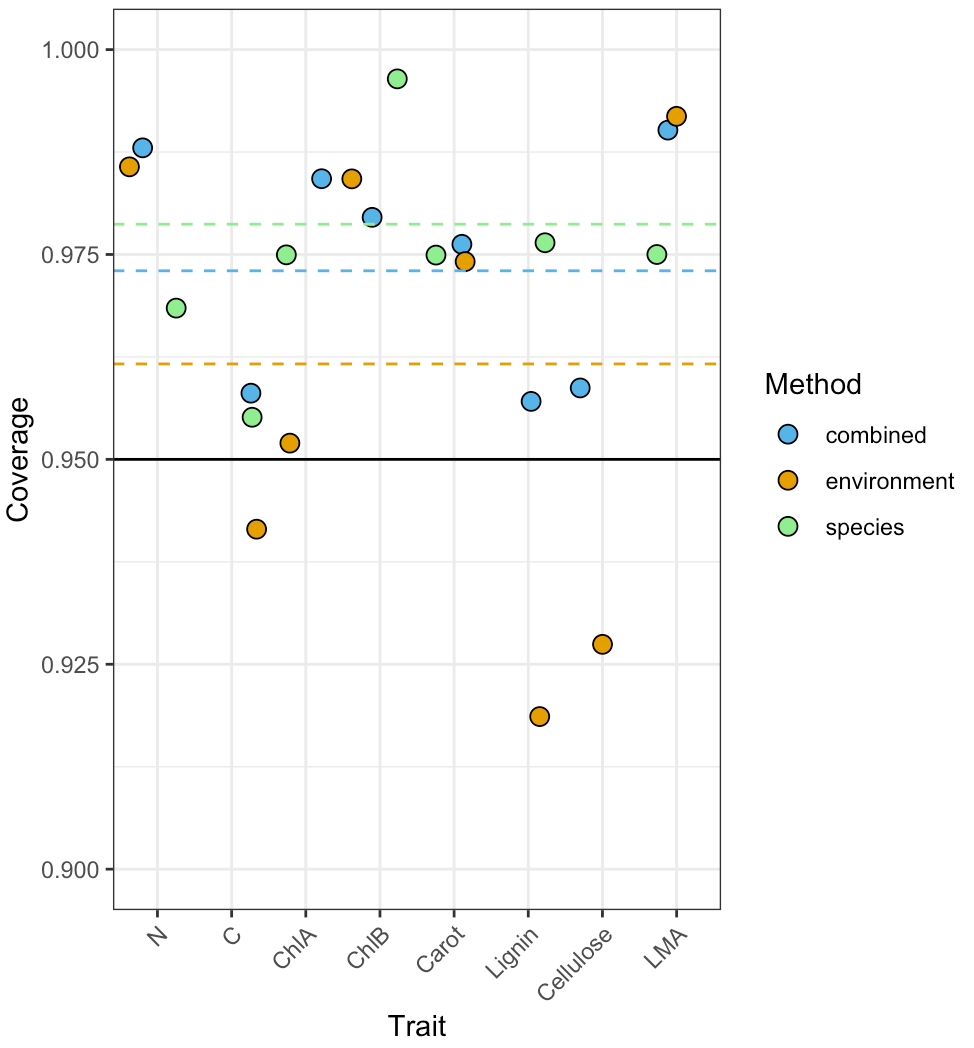

Figure S.4. Assessment of uncertainty estimates for the three models based on 95% coverage (the proportion of data points falling within the 95% prediction interval). The ideal value of 0.95 is indicated by the black solid line. Coverage for each trait is represented by orange dots (environmental-only model), green (phylogeny-only model) and blue (combined model). Average coverage for the three models is indicated by dashed lines following the same color scheme.

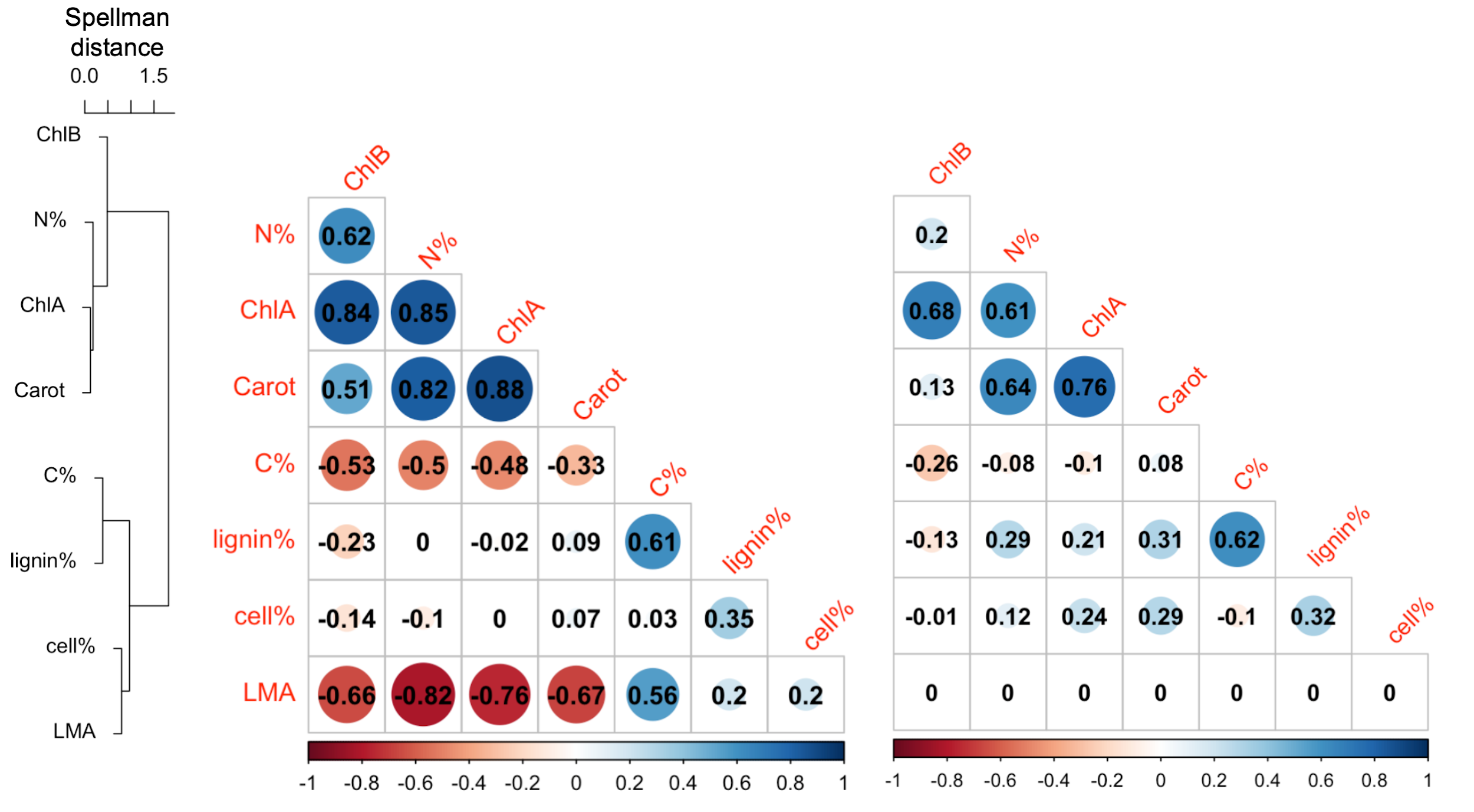

Figure S.5. Hierarchical clustering (left) and correlation matrix (right) based on residuals from the combined model for the 8 foliar traits included in this study. Model residuals represent variation in leaf traits not explained by any of the features included in the model. The residuals correlation matrix is therefore an estimate of ‘inherent’ trait correlations; i.e., trade-offs and correlations that occur within species in a given environment. Hierarchical clustering was based on complete linkage and shows that leaf traits included in this study can be grouped in two clusters: traits directly involved in photosynthesis (N% and pigments), and those involved in leaf structure (C%, lignin% and cellulose).

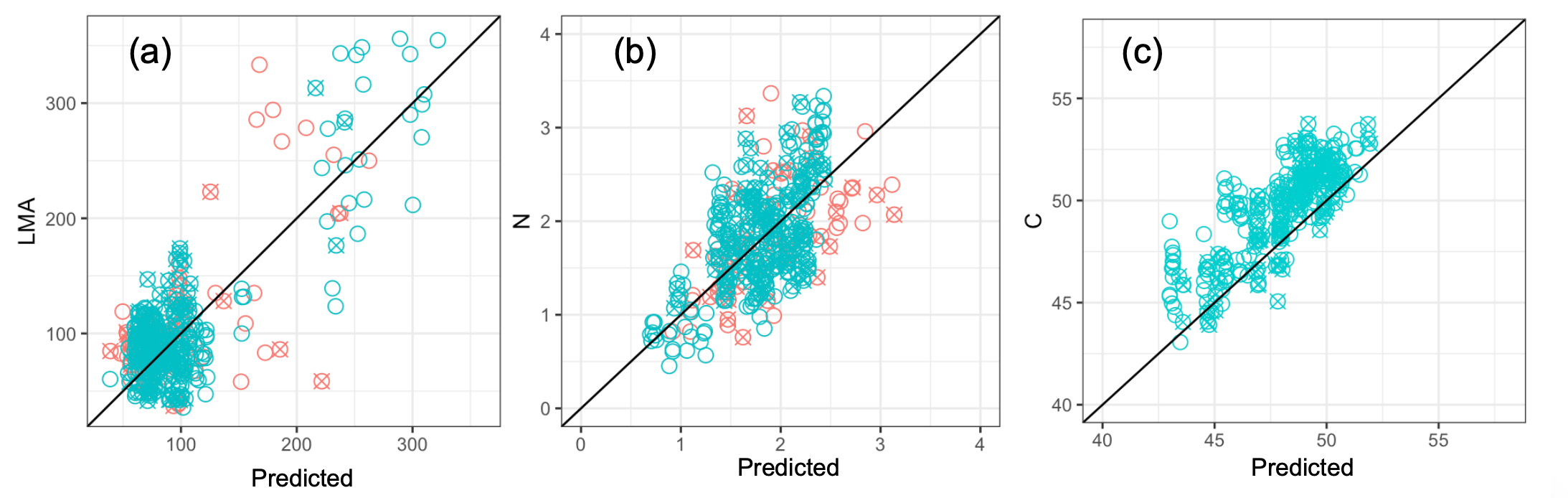

**Figure S.6. Transferability of the combined model to site and species outside of the training dataset for (a) LMA (g m^-2^), and (b) N%, and (c) C%. Filled dots are species present in the training set, hollow dots are species not trained on. Orange dots are individual trees from the BIEN dataset (n=88), and azure dots are from the TRY subset (n=265).**

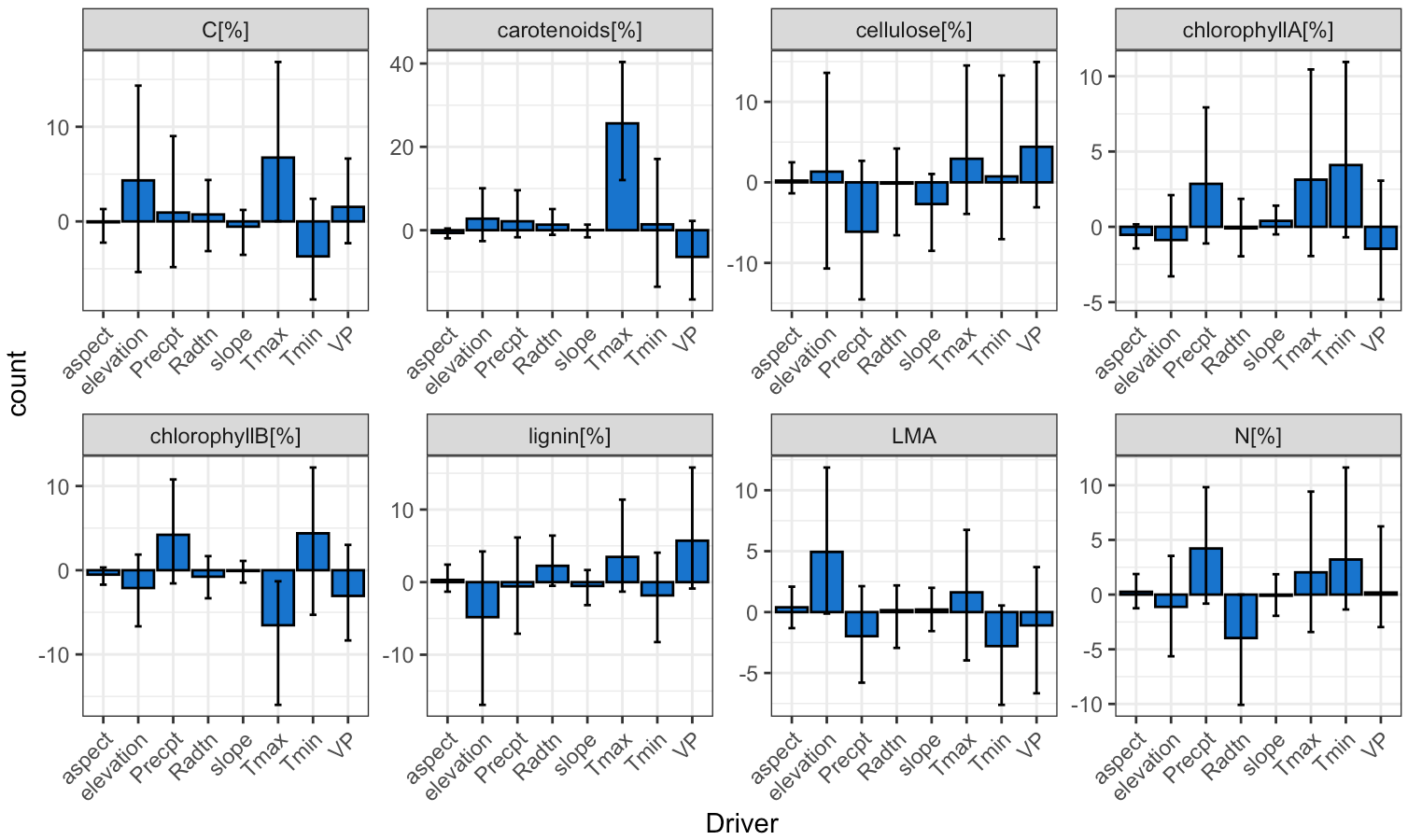

Figure S.7 Estimates of fixed effects coefficients for environmental variables calculated from sampling the posterior distribution of the combined model. Barplot height represents the mean, error bars represent 95 credible intervals (central 95 percent interval for the posterior distribution marginalized on each predictor). Topographic variables consist of: slope (degrees), aspect (degrees) and elevation (meters). Environmental variables consist of: shortwave radiation (Rdtn, W/m^2^), water vapor pressure (VP,Pa ), max temperature (Tmax, ℃), min temperature (Tmin, ℃), and precipitation (Precpt, mm). Fixed effects (parameters at the level of population) were standardized with mean in 0 and by their standard deviation.

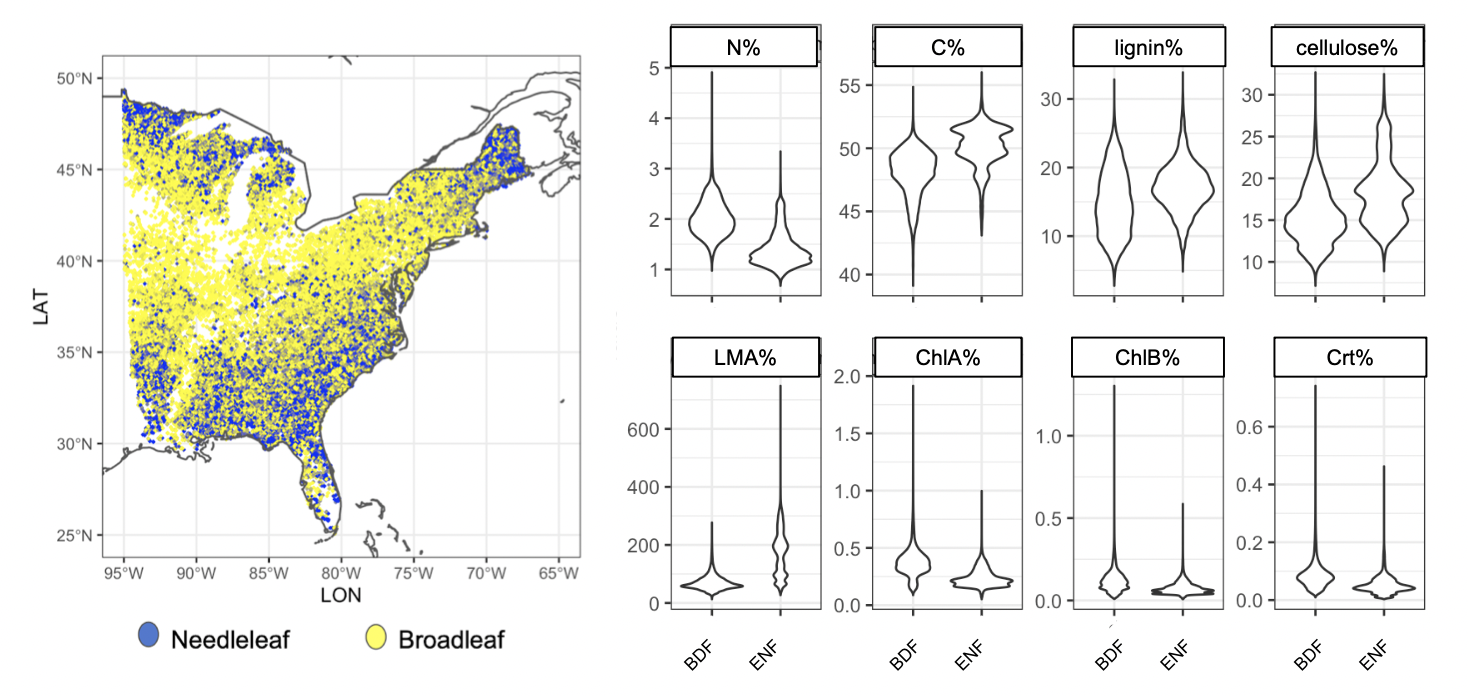

Figure S.8. Geographic distribution of FIA plots dominated by needleleaf species (blue) or broadleaf (yellow) species. “Dominated” here indicates that the functional group comprises >50% of abundance. Violin plots on the right show the different distribution in leaf traits for the two different functional groups.

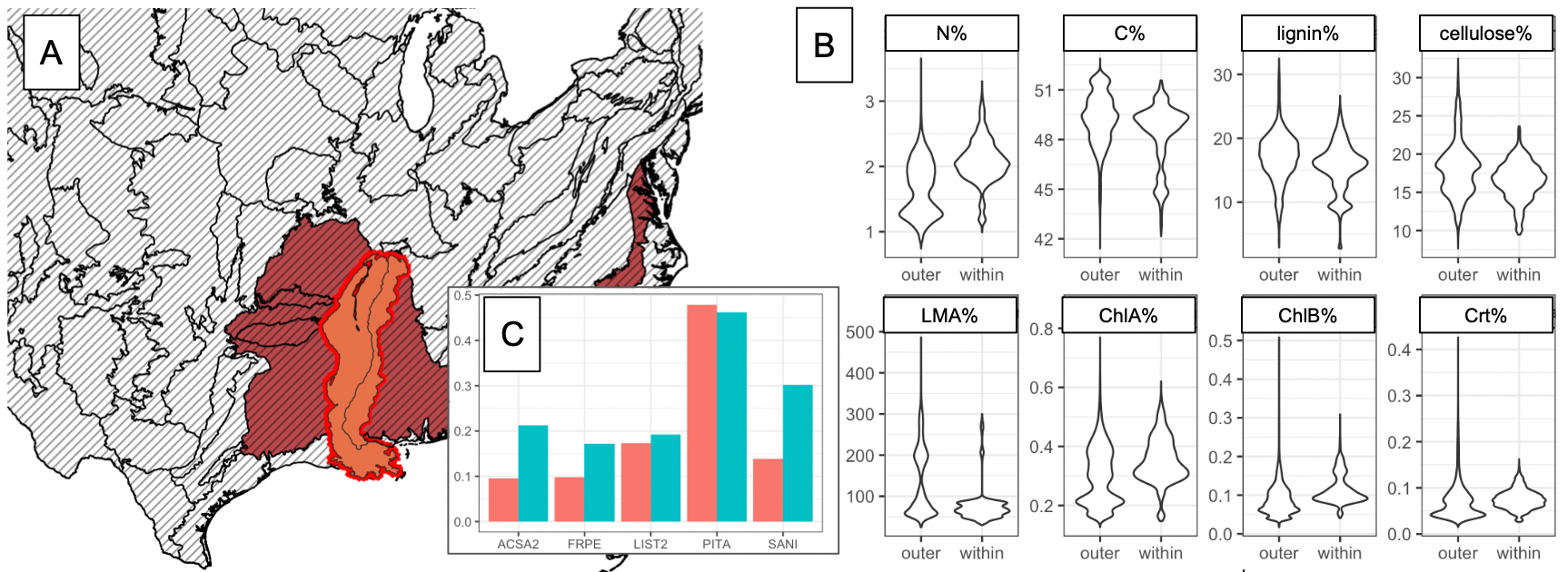

Figure S.9. (a) Geographic location of the Mississippi Alluvial Plains ecoregion (orange) and neighboring ecoregions (brown); (b) Violin plots show differences in leaf trait distributions for FIA plots in the Mississippi Alluvial Plains L3 ecoregion (labeled as “within”), and the other neighboring L3 ecoregions (“outer”); (c) relative abundance (number of trees over the total) of the 5 most abundant species in the Mississippi Alluvial Plains L3 ecoregion (blue) and in the neighboring L3 ecoregions (red). Although *Pinus palustris* (PIPA) is the most dominant species in both geographic areas, the Mississippi Alluvial Plains has higher abundance of *Fraxinus pennsylvanica* (FRPE), *Salix nigra* (SANI) and *Acer saccharum* (ACSA2); *Liquidambar styraciflua* (LIST2) is roughly equally abundant in the two areas.

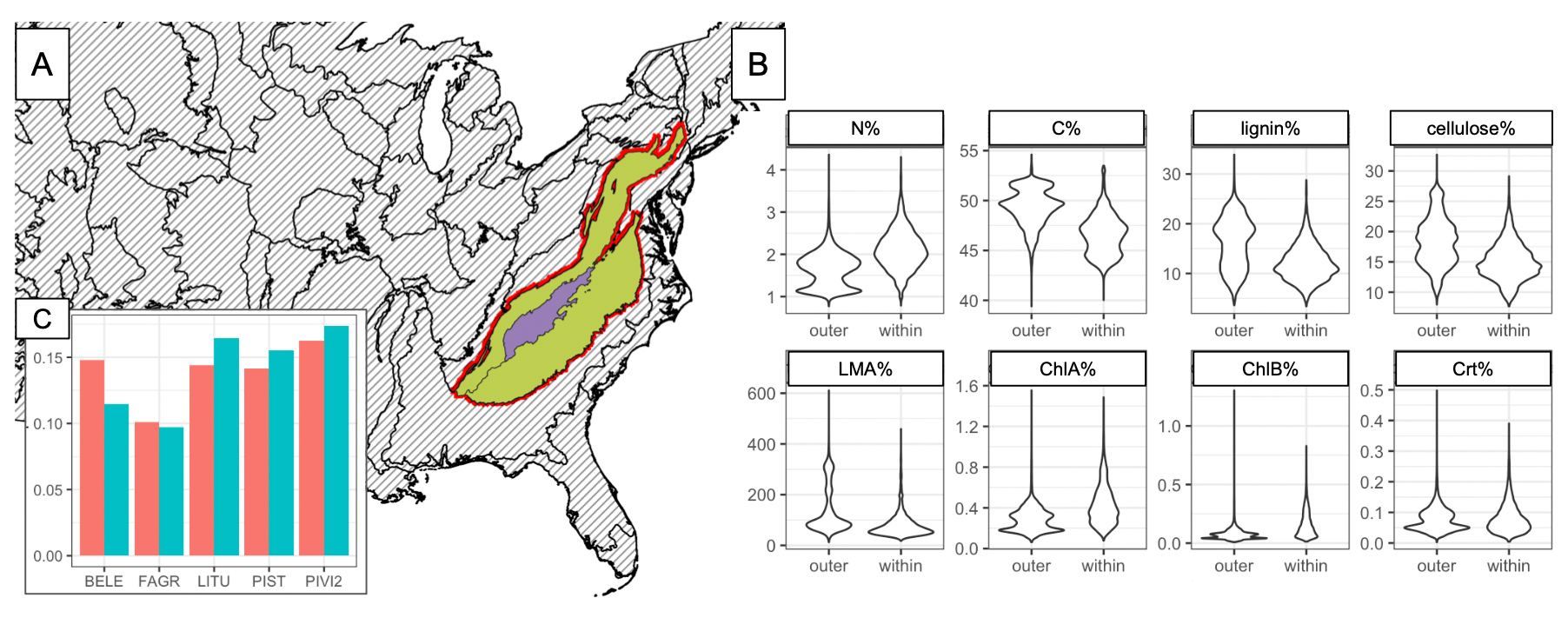

Figure S.10. (a) Geographic location of the Blue Ridge ecoregion (Purple) and neighboring ecoregions (Green); (b) Violin plots show differences in leaf trait distributions for FIA plots in the Blue Ridge L3 ecoregion (labeled as “within”), and the other neighboring L3 ecoregions (outer); (c) abundance of the 5 most abundant species in the Blue Ridge L3 ecoregion (blue) and in the other neighboring L3 ecoregions (red). According to the FIA census (2015-2019), all these regions are characterized by mixed vegetation, with dominant species including evergreen pines (mostly *Pinus strobus* and *Pinus virginiana*) and broadleaf species (mostly *Fagus grandifolia* and *Betula lenta*).

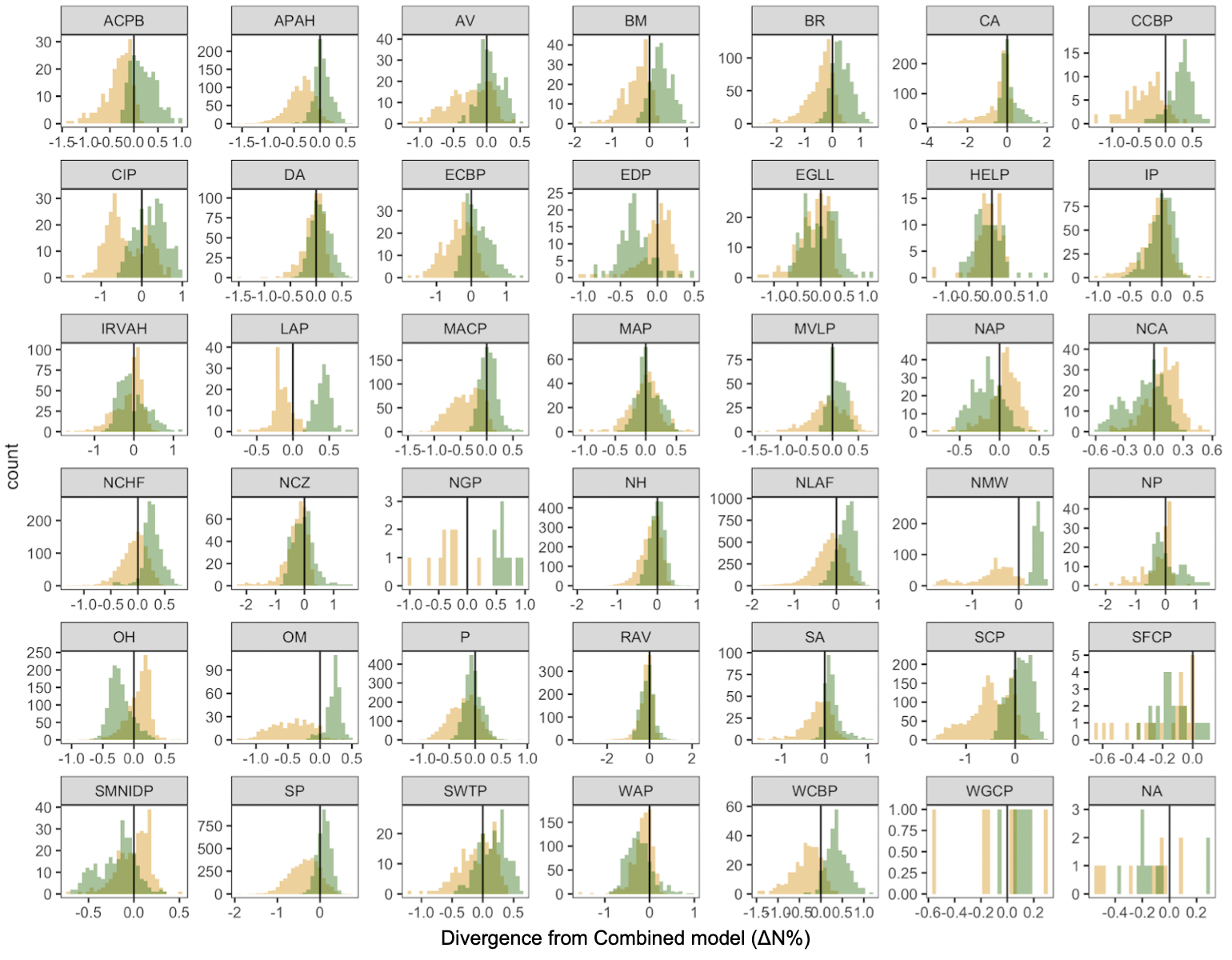

Figure S.11. (A) Distributions of N% divergence between the combined model and the environmental model (yellow) or the species model (green) for each L3 ecoregion. Vertical black line represents FIA plots where the divergence from the combined model is 0. Ecoregions code names are: Southern Florida Coastal Plain = SFCP, Southern Coastal Plain = SCP , Mississippi Alluvial Plain = MAP, Southeastern Plains = SP, Western Gulf Coastal Plain = WGCP, Mississippi Valley Loess Plains = MVLP, South Central Plains = SCP, Middle Atlantic Coastal Plain = MACP, Piedmont = P, Ridge and Valley = RAV , Southwestern Appalachians = SA, Ouachita Mountains = OM, Blue Ridge = BR , Interior Plateau = IP, Arkansas Valley = AV, Boston Mountains = BM, Ozark Highlands = OH, Central Appalachians = CA, Interior River Valleys and Hills = IRVAH, Western Allegheny Plateau = WAP, Central Irregular Plains = CIP, Northern Piedmont = NP, Eastern Corn Belt Plains = ECBP, Atlantic Coastal Pine Barrens = ACPB, Western Corn Belt Plains = WCBP, Central Corn Belt Plains = CCBP, Erie Drift Plain = EDP, Northeastern Highlands = NH, Northeastern Coastal Zone = NCZ, Southern Michigan/Northern Indiana Drift Plains = SMIDP, North Central Appalachians = NCA, Huron/Erie Lake Plains = HLP, Northern Allegheny Plateau = NAP, Eastern Great Lakes Lowlands = EGLL, Driftless Area = DA, Southeastern Wisconsin Till Plains = SWTP, Northern Lakes and Forests = NLAF, North Central Hardwood Forests = NCHF , Acadian Plains and Hills = APAH, Northern Glaciated Plains = NGP, Lake Agassiz Plain = LAP, Northern Minnesota Wetlands = NMW.

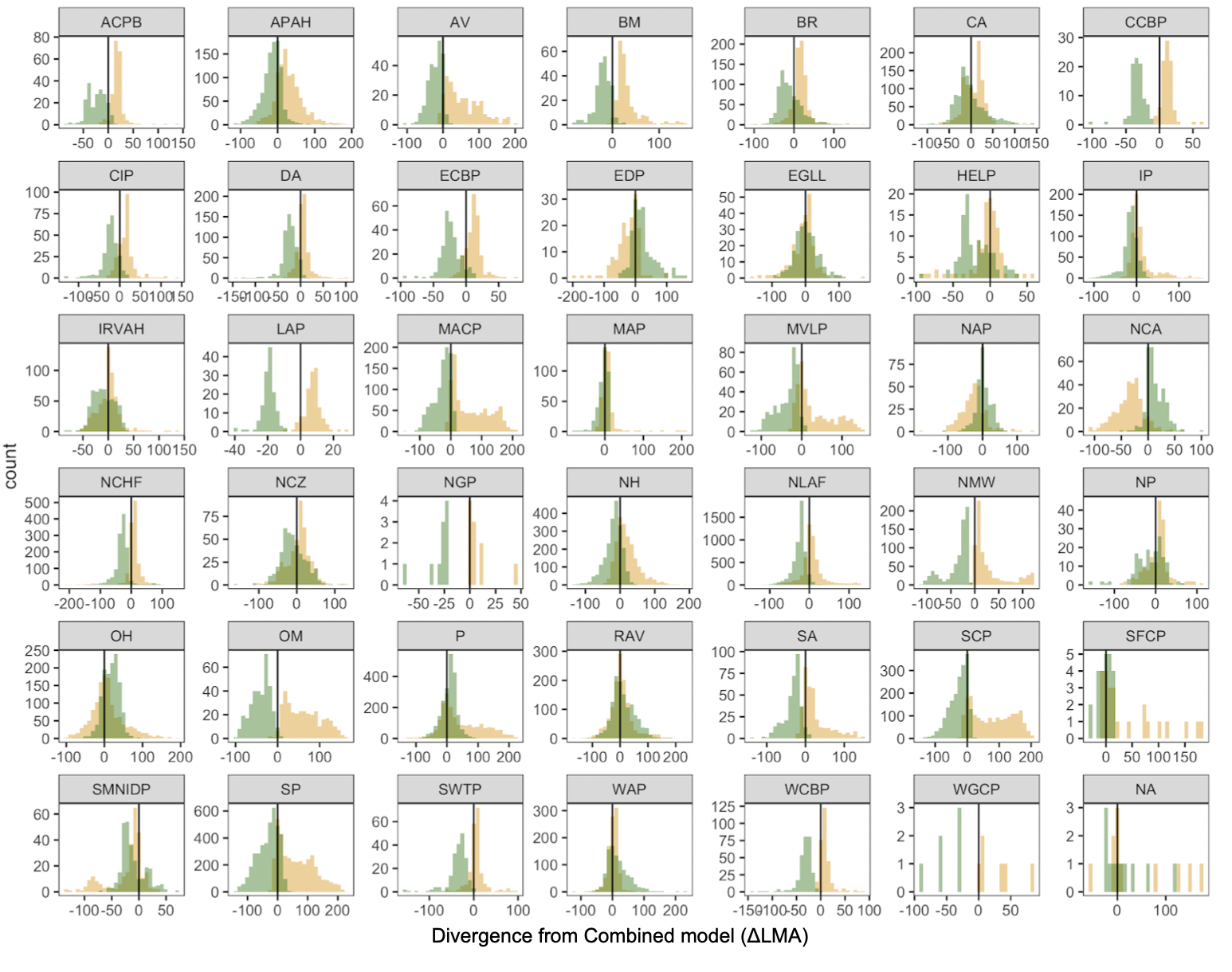
Figure S.11 (continued). (B) Distributions of LMA divergence between the combined model and the environmental model (yellow) or the species model (green) for each L3 ecoregion. Vertical black line represents FIA plots where the divergence from the combined model is 0. Ecoregions code names are: Southern Florida Coastal Plain = SFCP, Southern Coastal Plain = SCP , Mississippi Alluvial Plain = MAP, Southeastern Plains = SP, Western Gulf Coastal Plain = WGCP, Mississippi Valley Loess Plains = MVLP, South Central Plains = SCP, Middle Atlantic Coastal Plain = MACP, Piedmont = P, Ridge and Valley = RAV , Southwestern Appalachians = SA, Ouachita Mountains = OM, Blue Ridge = BR , Interior Plateau = IP, Arkansas Valley = AV, Boston Mountains = BM, Ozark Highlands = OH, Central Appalachians = CA, Interior River Valleys and Hills = IRVAH, Western Allegheny Plateau = WAP, Central Irregular Plains = CIP, Northern Piedmont = NP, Eastern Corn Belt Plains = ECBP, Atlantic Coastal Pine Barrens = ACPB, Western Corn Belt Plains = WCBP, Central Corn Belt Plains = CCBP, Erie Drift Plain = EDP, Northeastern Highlands = NH, Northeastern Coastal Zone = NCZ, Southern Michigan/Northern Indiana Drift Plains = SMIDP, North Central Appalachians = NCA, Huron/Erie Lake Plains = HLP, Northern Allegheny Plateau = NAP, Eastern Great Lakes Lowlands = EGLL, Driftless Area = DA, Southeastern Wisconsin Till Plains = SWTP, Northern Lakes and Forests = NLAF, North Central Hardwood Forests = NCHF , Acadian Plains and Hills = APAH, Northern Glaciated Plains = NGP, Lake Agassiz Plain = LAP, Northern Minnesota Wetlands = NMW.

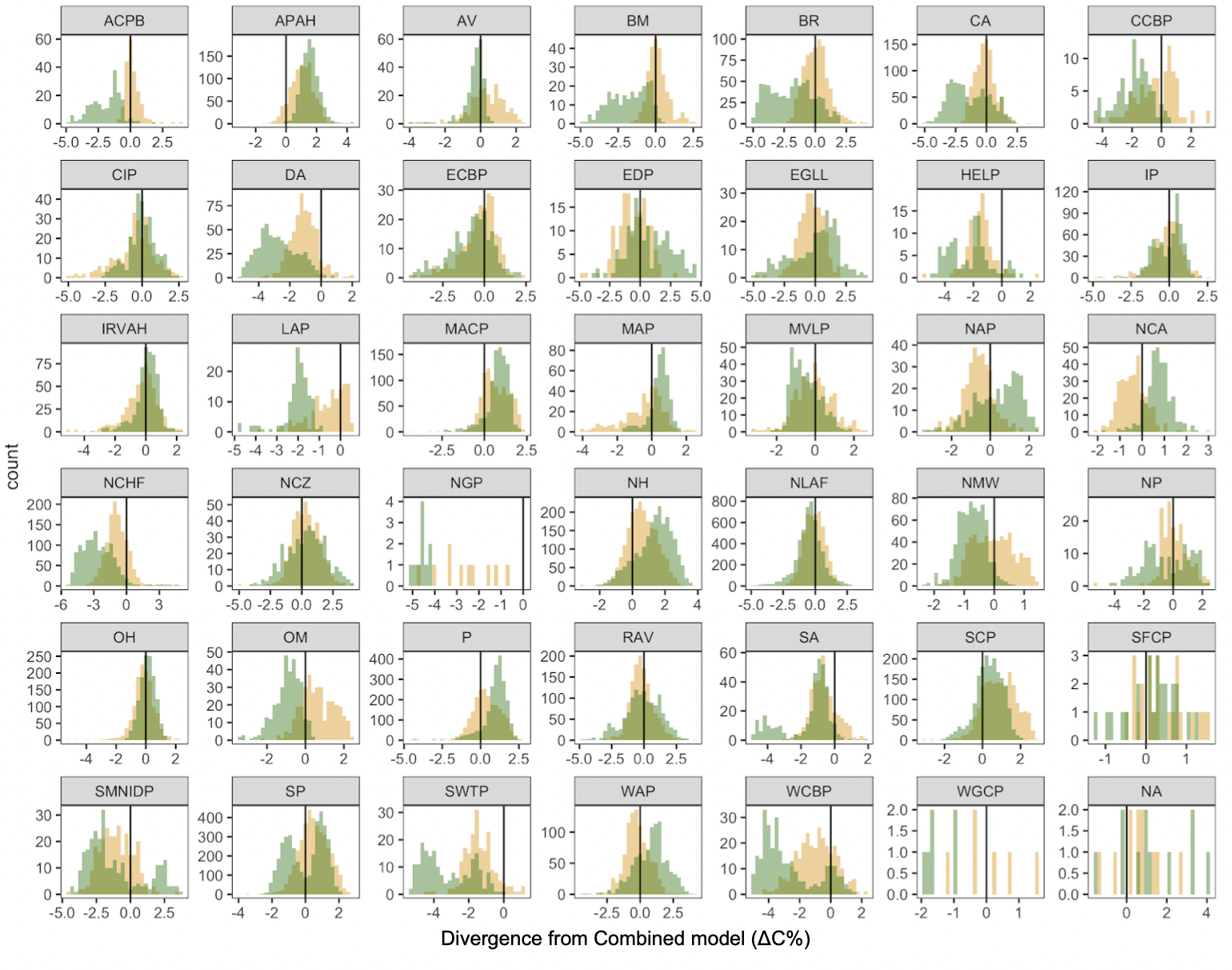
Figure S.11 (continued). (C) Distributions of C% divergence between the combined model and the environmental model (yellow) or the species model (green) for each L3 ecoregion. Vertical black line represents FIA plots where the divergence from the combined model is 0. Ecoregions code names are: Southern Florida Coastal Plain = SFCP, Southern Coastal Plain = SCP , Mississippi Alluvial Plain = MAP, Southeastern Plains = SP, Western Gulf Coastal Plain = WGCP, Mississippi Valley Loess Plains = MVLP, South Central Plains = SCP, Middle Atlantic Coastal Plain = MACP, Piedmont = P, Ridge and Valley = RAV , Southwestern Appalachians = SA, Ouachita Mountains = OM, Blue Ridge = BR , Interior Plateau = IP, Arkansas Valley = AV, Boston Mountains = BM, Ozark Highlands = OH, Central Appalachians = CA, Interior River Valleys and Hills = IRVAH, Western Allegheny Plateau = WAP, Central Irregular Plains = CIP, Northern Piedmont = NP, Eastern Corn Belt Plains = ECBP, Atlantic Coastal Pine Barrens = ACPB, Western Corn Belt Plains = WCBP, Central Corn Belt Plains = CCBP, Erie Drift Plain = EDP, Northeastern Highlands = NH, Northeastern Coastal Zone = NCZ, Southern Michigan/Northern Indiana Drift Plains = SMIDP, North Central Appalachians = NCA, Huron/Erie Lake Plains = HLP, Northern Allegheny Plateau = NAP, Eastern Great Lakes Lowlands = EGLL, Driftless Area = DA, Southeastern Wisconsin Till Plains = SWTP, Northern Lakes and Forests = NLAF, North Central Hardwood Forests = NCHF , Acadian Plains and Hills = APAH, Northern Glaciated Plains = NGP, Lake Agassiz Plain = LAP, Northern Minnesota Wetlands = NMW.

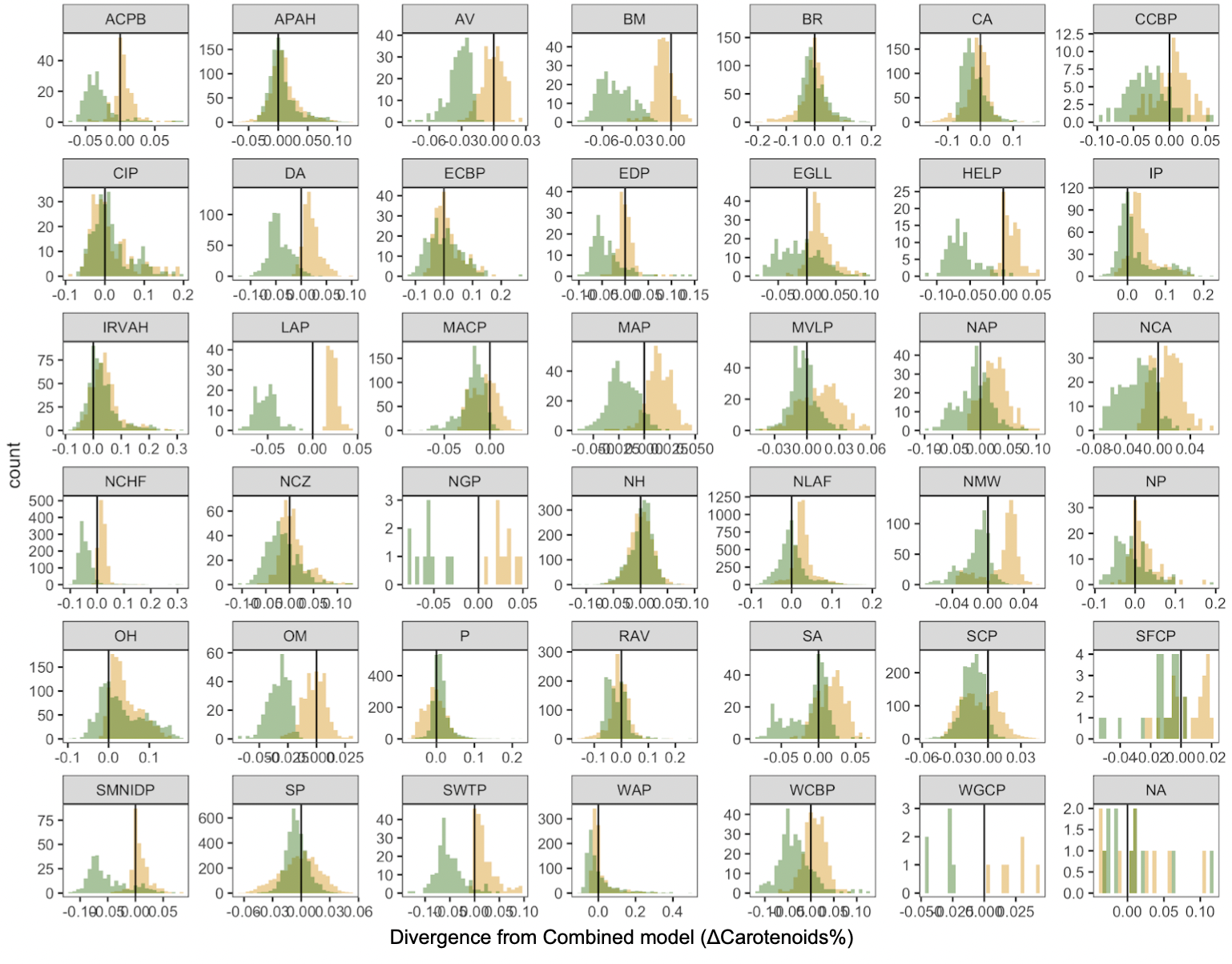
Figure S.11 (continued). (D) Distributions of carotenoids% divergence between the combined model and the environmental model (yellow) or the species model (green) for each L3 ecoregion. Vertical black line represents FIA plots where the divergence from the combined model is 0. Ecoregions code names are: Southern Florida Coastal Plain = SFCP, Southern Coastal Plain = SCP , Mississippi Alluvial Plain = MAP, Southeastern Plains = SP, Western Gulf Coastal Plain = WGCP, Mississippi Valley Loess Plains = MVLP, South Central Plains = SCP, Middle Atlantic Coastal Plain = MACP, Piedmont = P, Ridge and Valley = RAV , Southwestern Appalachians = SA, Ouachita Mountains = OM, Blue Ridge = BR , Interior Plateau = IP, Arkansas Valley = AV, Boston Mountains = BM, Ozark Highlands = OH, Central Appalachians = CA, Interior River Valleys and Hills = IRVAH, Western Allegheny Plateau = WAP, Central Irregular Plains = CIP, Northern Piedmont = NP, Eastern Corn Belt Plains = ECBP, Atlantic Coastal Pine Barrens = ACPB, Western Corn Belt Plains = WCBP, Central Corn Belt Plains = CCBP, Erie Drift Plain = EDP, Northeastern Highlands = NH, Northeastern Coastal Zone = NCZ, Southern Michigan/Northern Indiana Drift Plains = SMIDP, North Central Appalachians = NCA, Huron/Erie Lake Plains = HLP, Northern Allegheny Plateau = NAP, Eastern Great Lakes Lowlands = EGLL, Driftless Area = DA, Southeastern Wisconsin Till Plains = SWTP, Northern Lakes and Forests = NLAF, North Central Hardwood Forests = NCHF , Acadian Plains and Hills = APAH, Northern Glaciated Plains = NGP, Lake Agassiz Plain = LAP, Northern Minnesota Wetlands = NMW.

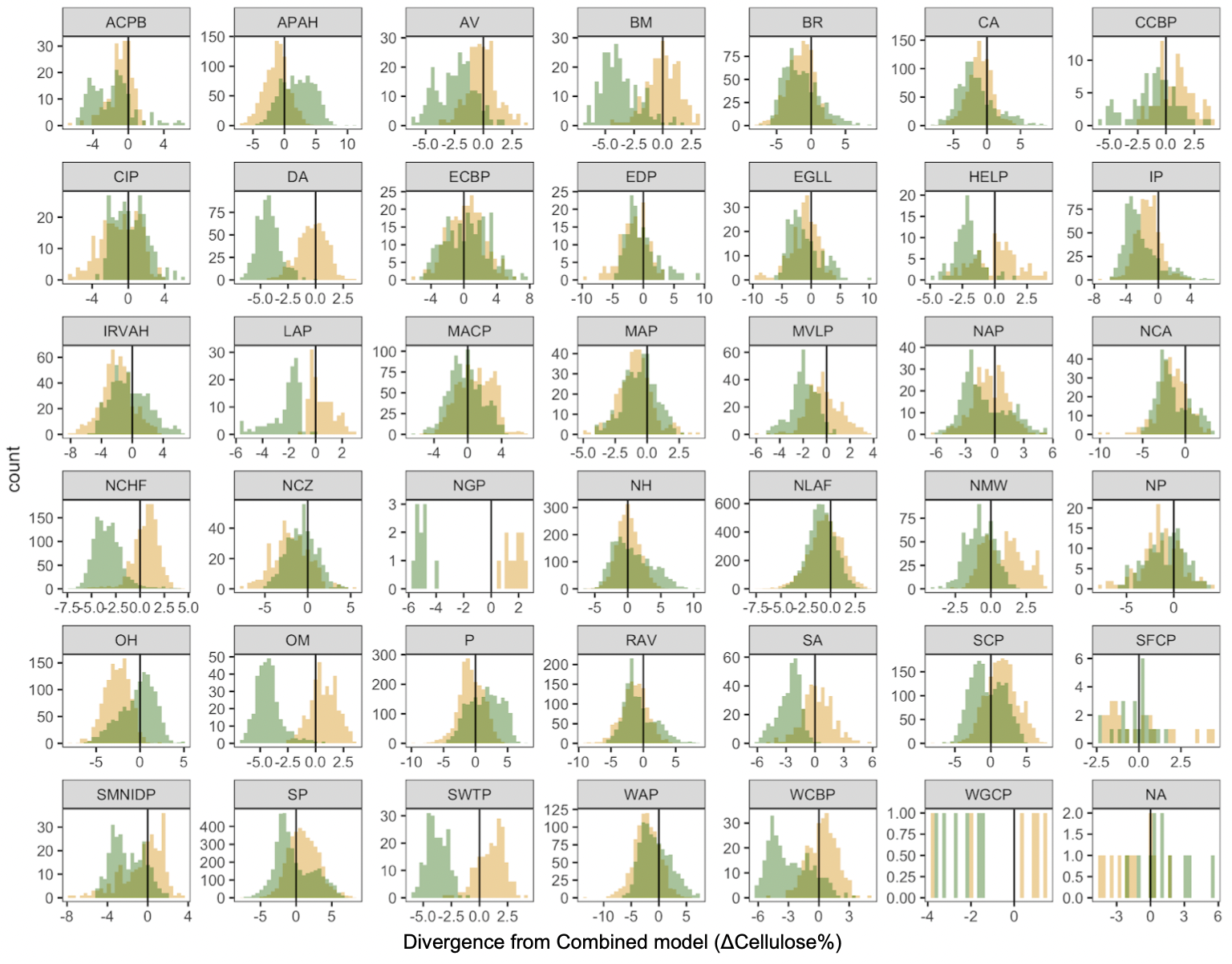
Figure S.11 (continued). (E) Distributions of cellulose% divergence between the combined model and the environmental model (yellow) or the species model (green) for each L3 ecoregion. Vertical black line represents FIA plots where the divergence from the combined model is 0. Ecoregions code names are: Southern Florida Coastal Plain = SFCP, Southern Coastal Plain = SCP , Mississippi Alluvial Plain = MAP, Southeastern Plains = SP, Western Gulf Coastal Plain = WGCP, Mississippi Valley Loess Plains = MVLP, South Central Plains = SCP, Middle Atlantic Coastal Plain = MACP, Piedmont = P, Ridge and Valley = RAV , Southwestern Appalachians = SA, Ouachita Mountains = OM, Blue Ridge = BR , Interior Plateau = IP, Arkansas Valley = AV, Boston Mountains = BM, Ozark Highlands = OH, Central Appalachians = CA, Interior River Valleys and Hills = IRVAH, Western Allegheny Plateau = WAP, Central Irregular Plains = CIP, Northern Piedmont = NP, Eastern Corn Belt Plains = ECBP, Atlantic Coastal Pine Barrens = ACPB, Western Corn Belt Plains = WCBP, Central Corn Belt Plains = CCBP, Erie Drift Plain = EDP, Northeastern Highlands = NH, Northeastern Coastal Zone = NCZ, Southern Michigan/Northern Indiana Drift Plains = SMIDP, North Central Appalachians = NCA, Huron/Erie Lake Plains = HLP, Northern Allegheny Plateau = NAP, Eastern Great Lakes Lowlands = EGLL, Driftless Area = DA, Southeastern Wisconsin Till Plains = SWTP, Northern Lakes and Forests = NLAF, North Central Hardwood Forests = NCHF , Acadian Plains and Hills = APAH, Northern Glaciated Plains = NGP, Lake Agassiz Plain = LAP, Northern Minnesota Wetlands = NMW.

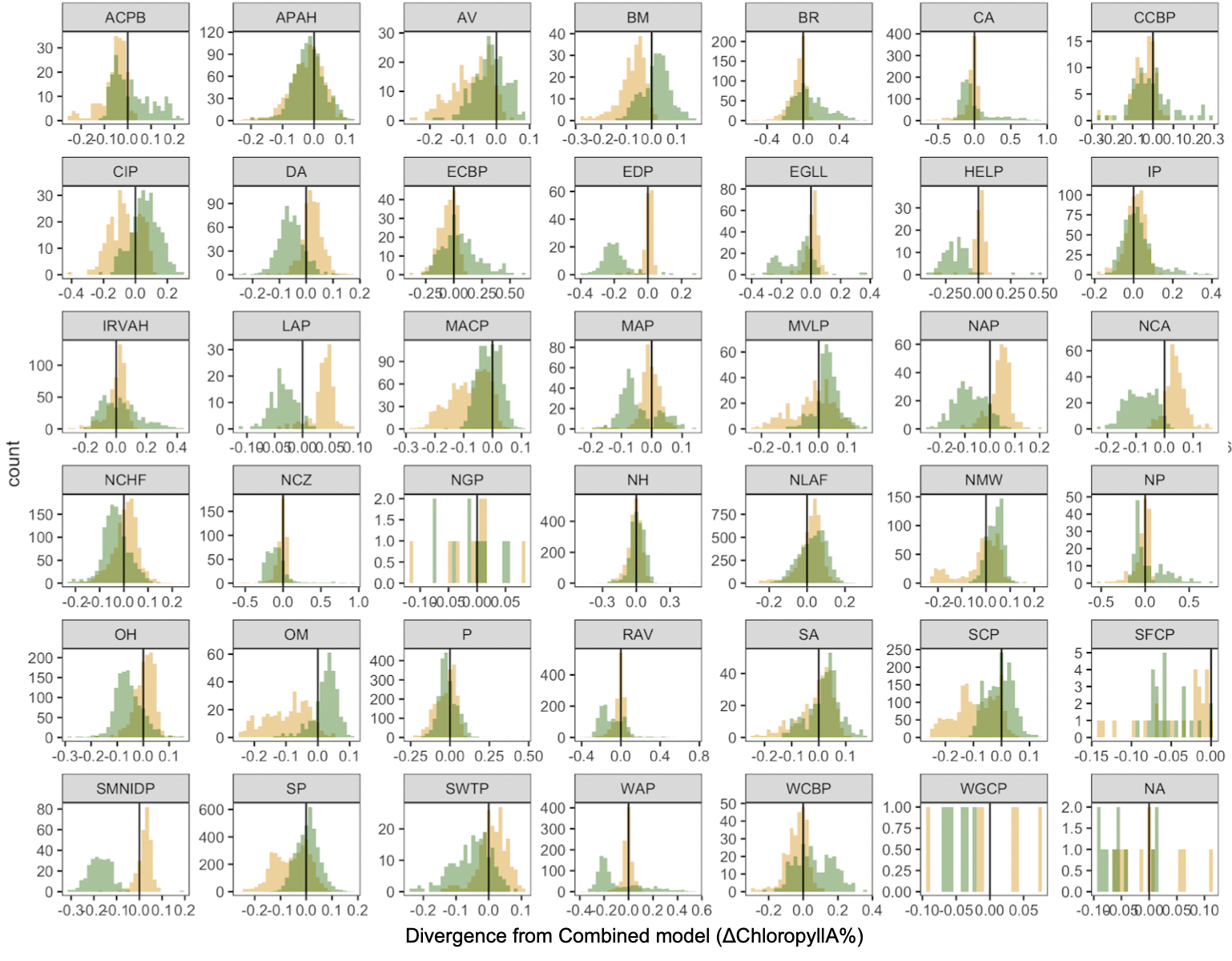

Figure S.11 (continued). (F) Distributions of chlorophyll A% divergence between the combined model and the environmental model (yellow) or the species model (green) for each L3 ecoregion. Vertical black line represents FIA plots where the divergence from the combined model is 0.Ecoregions code names are: Southern Florida Coastal Plain = SFCP, Southern Coastal Plain = SCP , Mississippi Alluvial Plain = MAP, Southeastern Plains = SP, Western Gulf Coastal Plain = WGCP, Mississippi Valley Loess Plains = MVLP, South Central Plains = SCP, Middle Atlantic Coastal Plain = MACP, Piedmont = P, Ridge and Valley = RAV , Southwestern Appalachians = SA, Ouachita Mountains = OM, Blue Ridge = BR , Interior Plateau = IP, Arkansas Valley = AV, Boston Mountains = BM, Ozark Highlands = OH, Central Appalachians = CA, Interior River Valleys and Hills = IRVAH, Western Allegheny Plateau = WAP, Central Irregular Plains = CIP, Northern Piedmont = NP, Eastern Corn Belt Plains = ECBP, Atlantic Coastal Pine Barrens = ACPB, Western Corn Belt Plains = WCBP, Central Corn Belt Plains = CCBP, Erie Drift Plain = EDP, Northeastern Highlands = NH, Northeastern Coastal Zone = NCZ, Southern Michigan/Northern Indiana Drift Plains = SMIDP, North Central Appalachians = NCA, Huron/Erie Lake Plains = HLP, Northern Allegheny Plateau = NAP, Eastern Great Lakes Lowlands = EGLL, Driftless Area = DA, Southeastern Wisconsin Till Plains = SWTP, Northern Lakes and Forests = NLAF, North Central Hardwood Forests = NCHF , Acadian Plains and Hills = APAH, Northern Glaciated Plains = NGP, Lake Agassiz Plain = LAP, Northern Minnesota Wetlands = NMW.

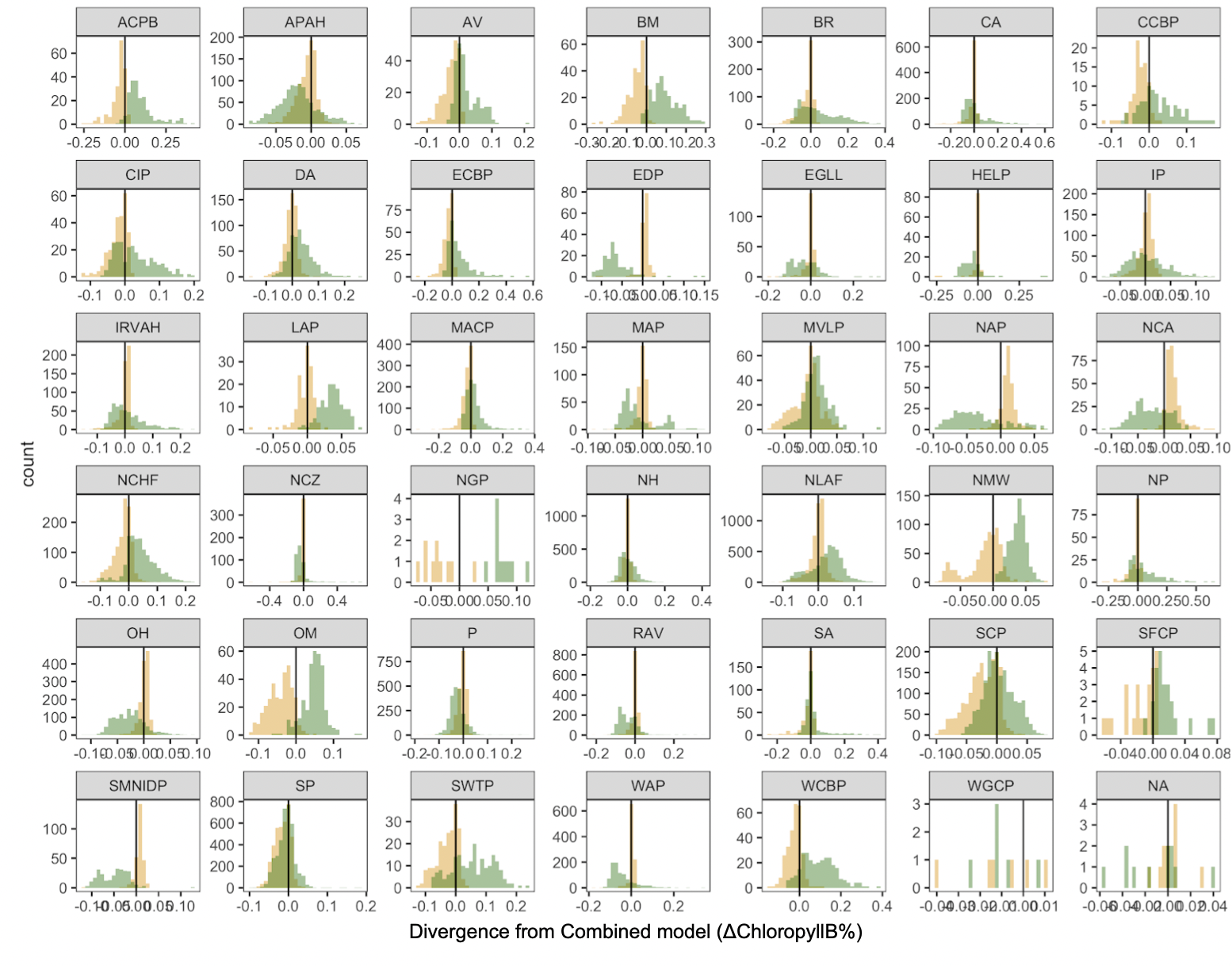
Figure S.11 (continued). (G) Distributions of chlorophyll B% divergence between the combined model and the environmental model (yellow) or the species model (green) for each L3 ecoregion. Vertical black line represents FIA plots where the divergence from the combined model is 0. Ecoregions code names are: Southern Florida Coastal Plain = SFCP, Southern Coastal Plain = SCP , Mississippi Alluvial Plain = MAP, Southeastern Plains = SP, Western Gulf Coastal Plain = WGCP, Mississippi Valley Loess Plains = MVLP, South Central Plains = SCP, Middle Atlantic Coastal Plain = MACP, Piedmont = P, Ridge and Valley = RAV , Southwestern Appalachians = SA, Ouachita Mountains = OM, Blue Ridge = BR , Interior Plateau = IP, Arkansas Valley = AV, Boston Mountains = BM, Ozark Highlands = OH, Central Appalachians = CA, Interior River Valleys and Hills = IRVAH, Western Allegheny Plateau = WAP, Central Irregular Plains = CIP, Northern Piedmont = NP, Eastern Corn Belt Plains = ECBP, Atlantic Coastal Pine Barrens = ACPB, Western Corn Belt Plains = WCBP, Central Corn Belt Plains = CCBP, Erie Drift Plain = EDP, Northeastern Highlands = NH, Northeastern Coastal Zone = NCZ, Southern Michigan/Northern Indiana Drift Plains = SMIDP, North Central Appalachians = NCA, Huron/Erie Lake Plains = HLP, Northern Allegheny Plateau = NAP, Eastern Great Lakes Lowlands = EGLL, Driftless Area = DA, Southeastern Wisconsin Till Plains = SWTP, Northern Lakes and Forests = NLAF, North Central Hardwood Forests = NCHF , Acadian Plains and Hills = APAH, Northern Glaciated Plains = NGP, Lake Agassiz Plain = LAP, Northern Minnesota Wetlands = NMW.
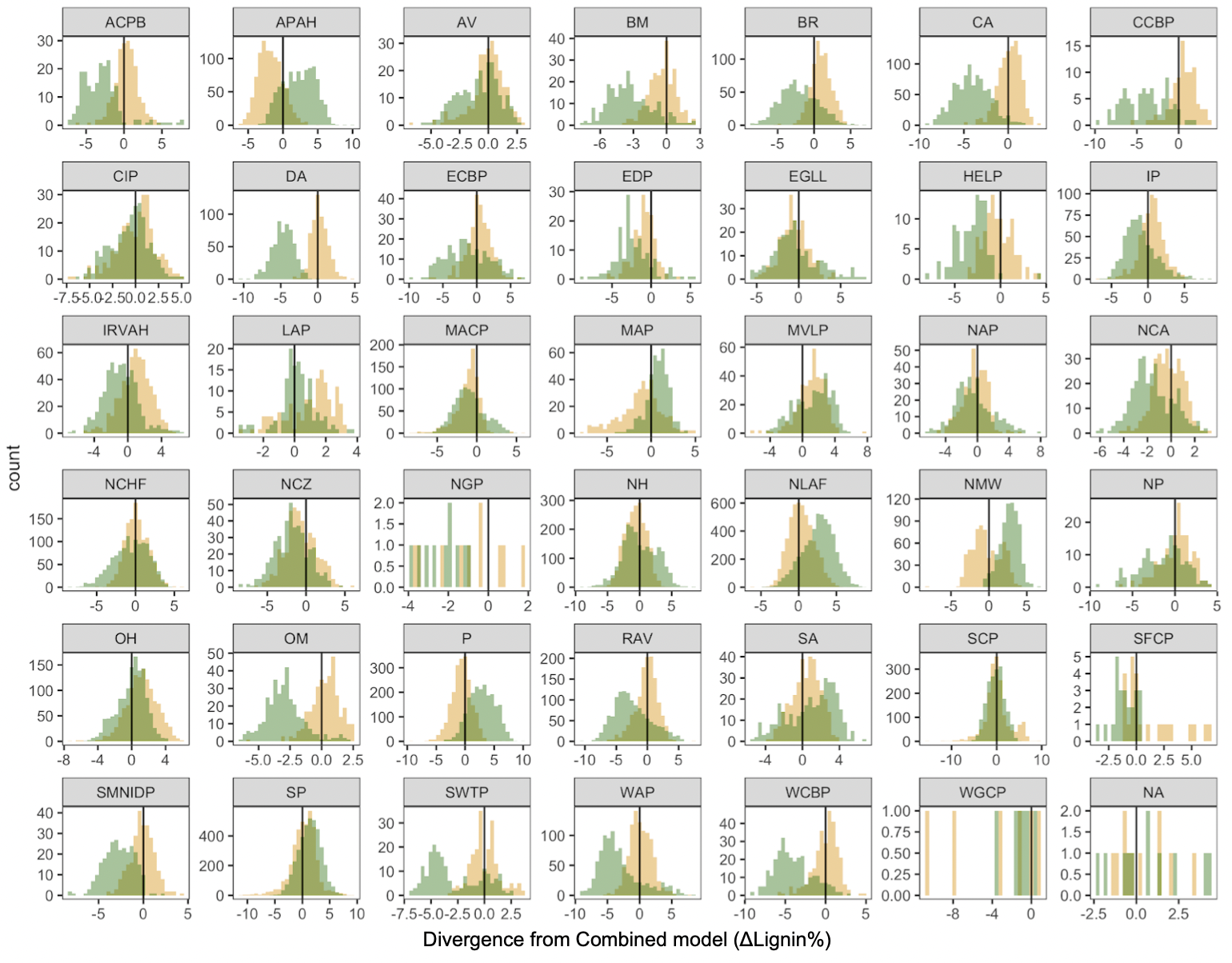
Figure S.11 (continued). (H) Distributions of Lignin% divergence between the combined model and the environmental model (yellow) or the species model (green) for each L3 ecoregion. Vertical black line represents FIA plots where the divergence from the combined model is 0. Ecoregions code names are: Southern Florida Coastal Plain = SFCP, Southern Coastal Plain = SCP , Mississippi Alluvial Plain = MAP, Southeastern Plains = SP, Western Gulf Coastal Plain = WGCP, Mississippi Valley Loess Plains = MVLP, South Central Plains = SCP, Middle Atlantic Coastal Plain = MACP, Piedmont = P, Ridge and Valley = RAV , Southwestern Appalachians = SA, Ouachita Mountains = OM, Blue Ridge = BR , Interior Plateau = IP, Arkansas Valley = AV, Boston Mountains = BM, Ozark Highlands = OH, Central Appalachians = CA, Interior River Valleys and Hills = IRVAH, Western Allegheny Plateau = WAP, Central Irregular Plains = CIP, Northern Piedmont = NP, Eastern Corn Belt Plains = ECBP, Atlantic Coastal Pine Barrens = ACPB, Western Corn Belt Plains = WCBP, Central Corn Belt Plains = CCBP, Erie Drift Plain = EDP, Northeastern Highlands = NH, Northeastern Coastal Zone = NCZ, Southern Michigan/Northern Indiana Drift Plains = SMIDP, North Central Appalachians = NCA, Huron/Erie Lake Plains = HLP, Northern Allegheny Plateau = NAP, Eastern Great Lakes Lowlands = EGLL, Driftless Area = DA, Southeastern Wisconsin Till Plains = SWTP, Northern Lakes and Forests = NLAF, North Central Hardwood Forests = NCHF , Acadian Plains and Hills = APAH, Northern Glaciated Plains = NGP, Lake Agassiz Plain = LAP, Northern Minnesota Wetlands = NMW.

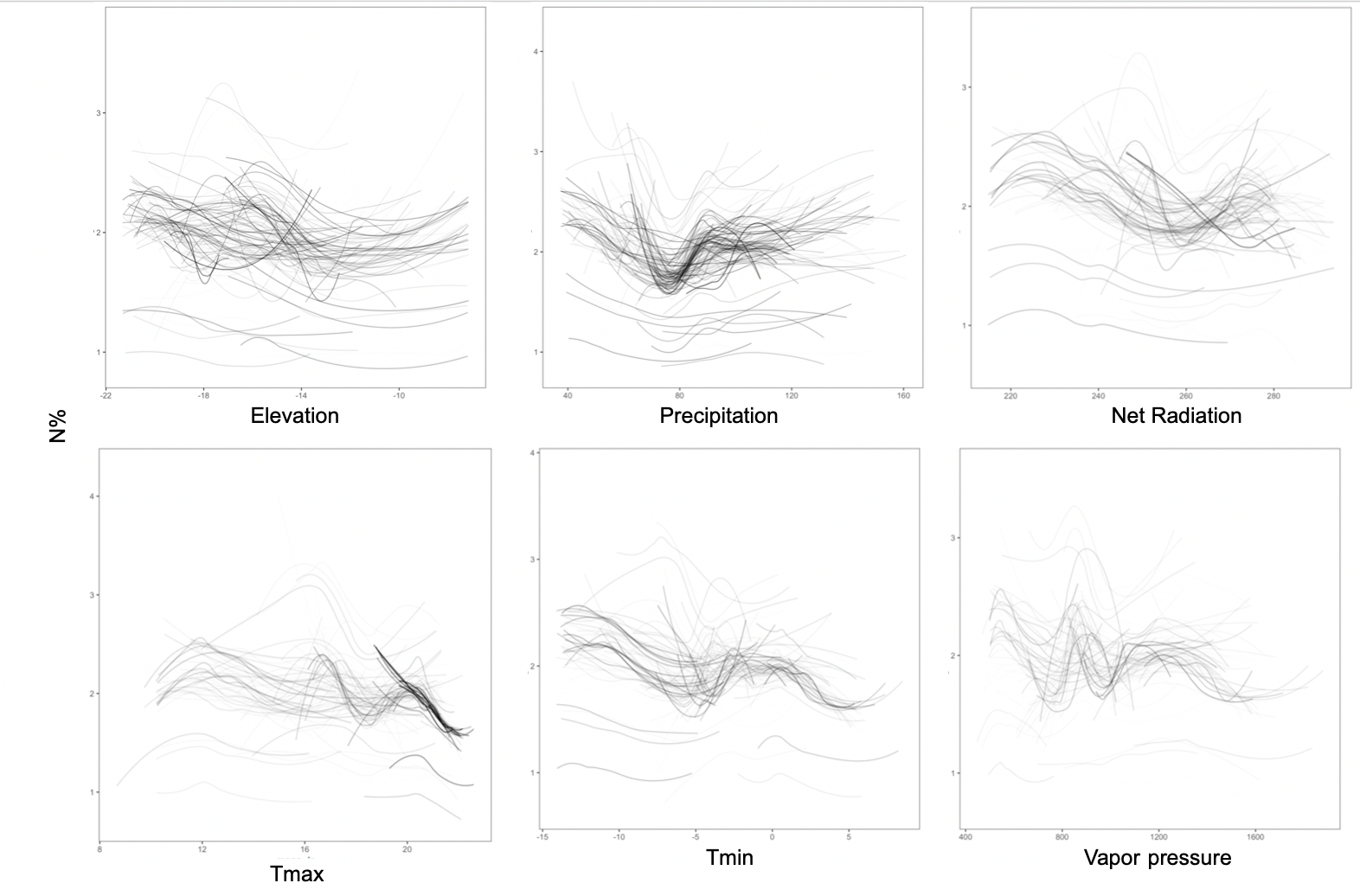

Figure S.12. Intraspecific variation of predicted leaf N% for tree species in eastern USA (200 species). Trends of the median N% prediction over the variation of environmental drivers (transparency matching the strength of the relationship for the species).

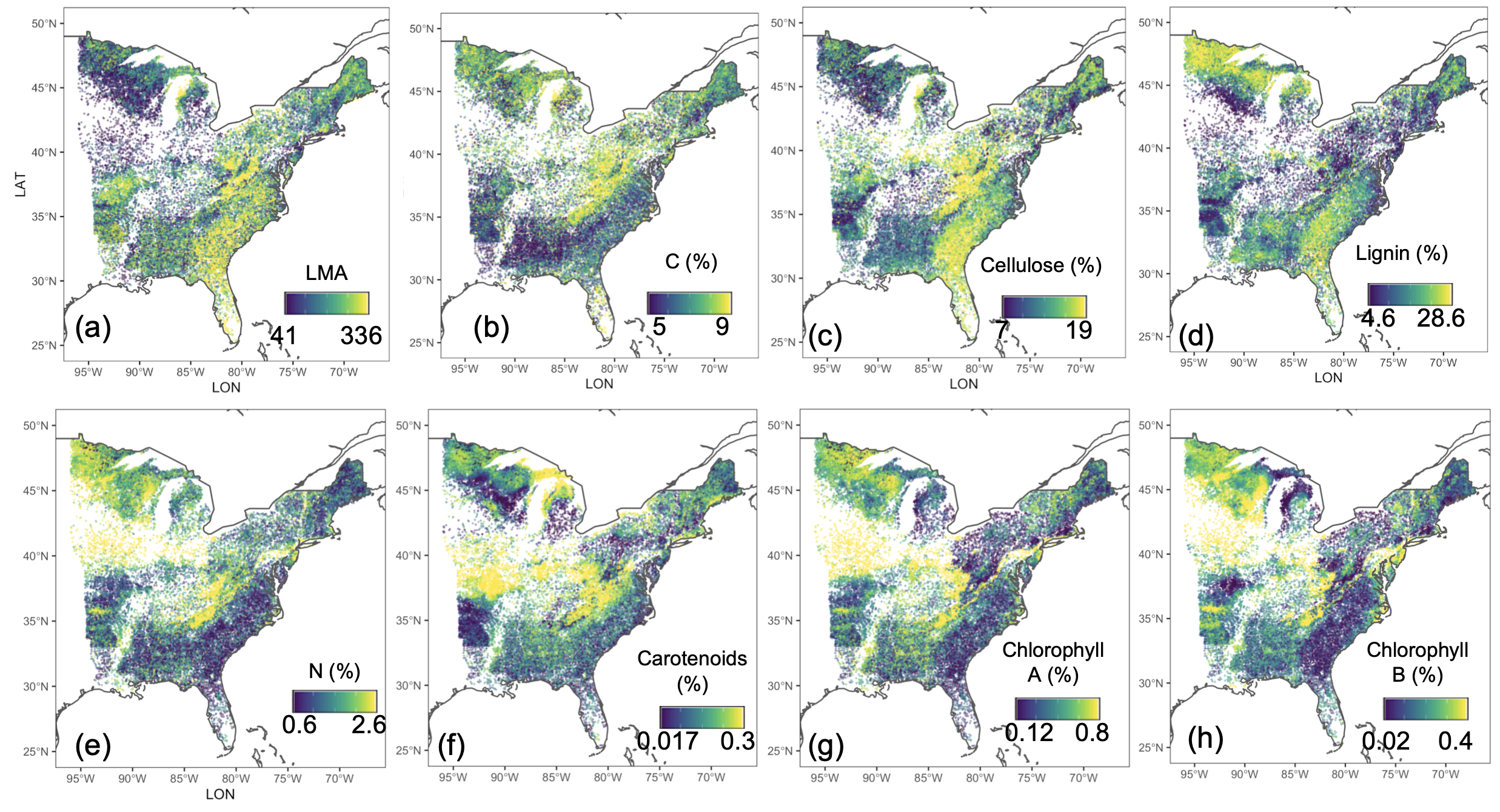

Figure S.13. Distribution of uncertainty from the combined model for the 8 leaf traits used in this study. Uncertainty (95PI range) was estimated for each individual tree in the FIA dataset and is shown here averaged across trees within FIA plots. Scale is from low ( blue) to high uncertainty (yellow). From top left to bottom right: chlorophyll A, chlorophyll B, carotenoids, N, C, lignin, cellulose, and LMA. Data to replicate and explore these patterns can be downloaded at https://doi.org/10.5281/zenodo.4647559.

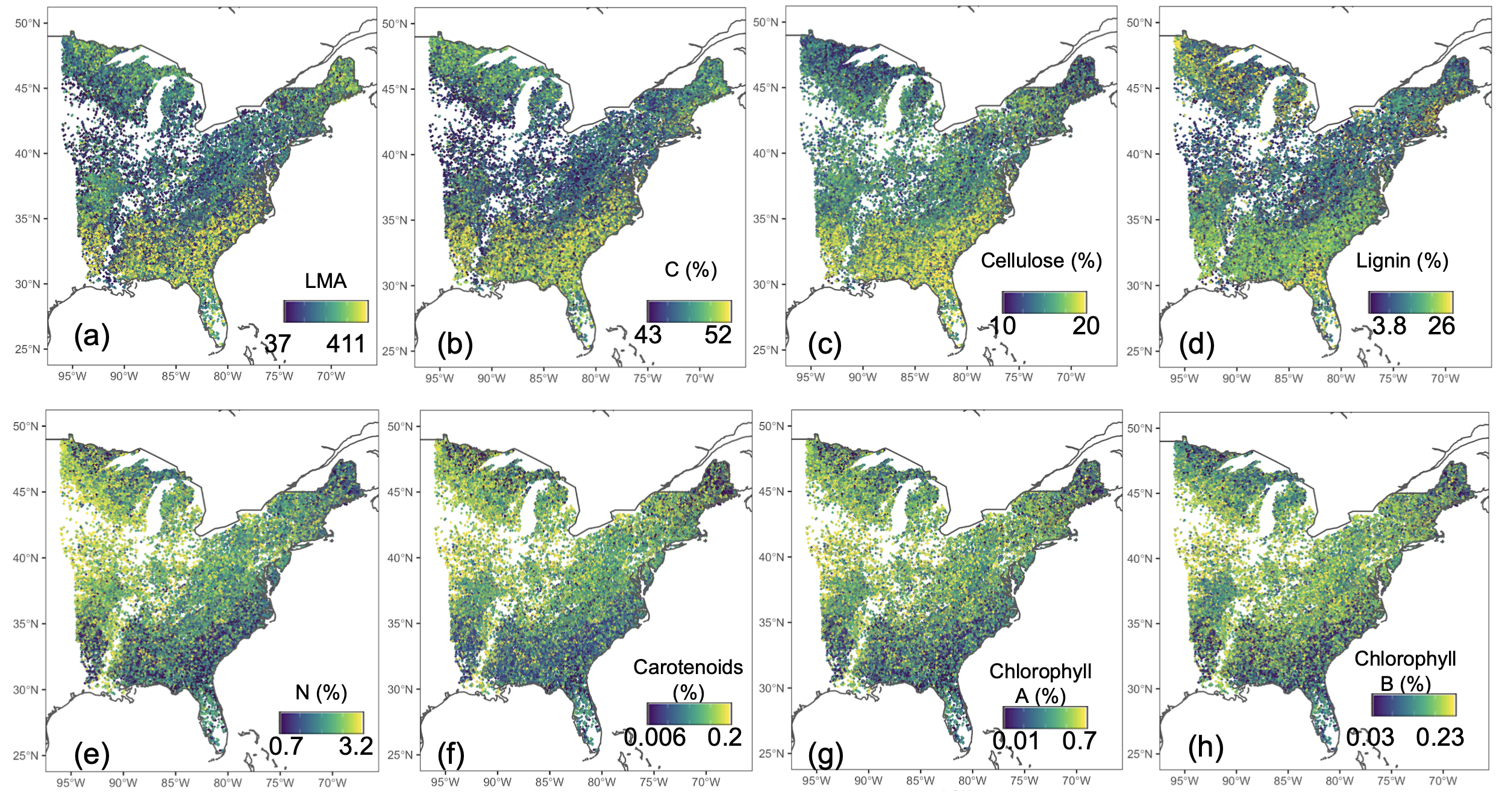

Figure S.13 (continued). Distribution of predictions from the phylogeny-only model for the 8 leaf traits used in this study. Predictions were estimated for each individual tree in the FIA dataset and are shown here averaged across trees within FIA plots. Scale is from low (blue) to high (yellow). From top left to bottom right: chlorophyll A, chlorophyll B, carotenoids, N, C, lignin, cellulose, and LMA. Data to replicate and explore these patterns can be downloaded at https://doi.org/10.5281/zenodo.4647559.

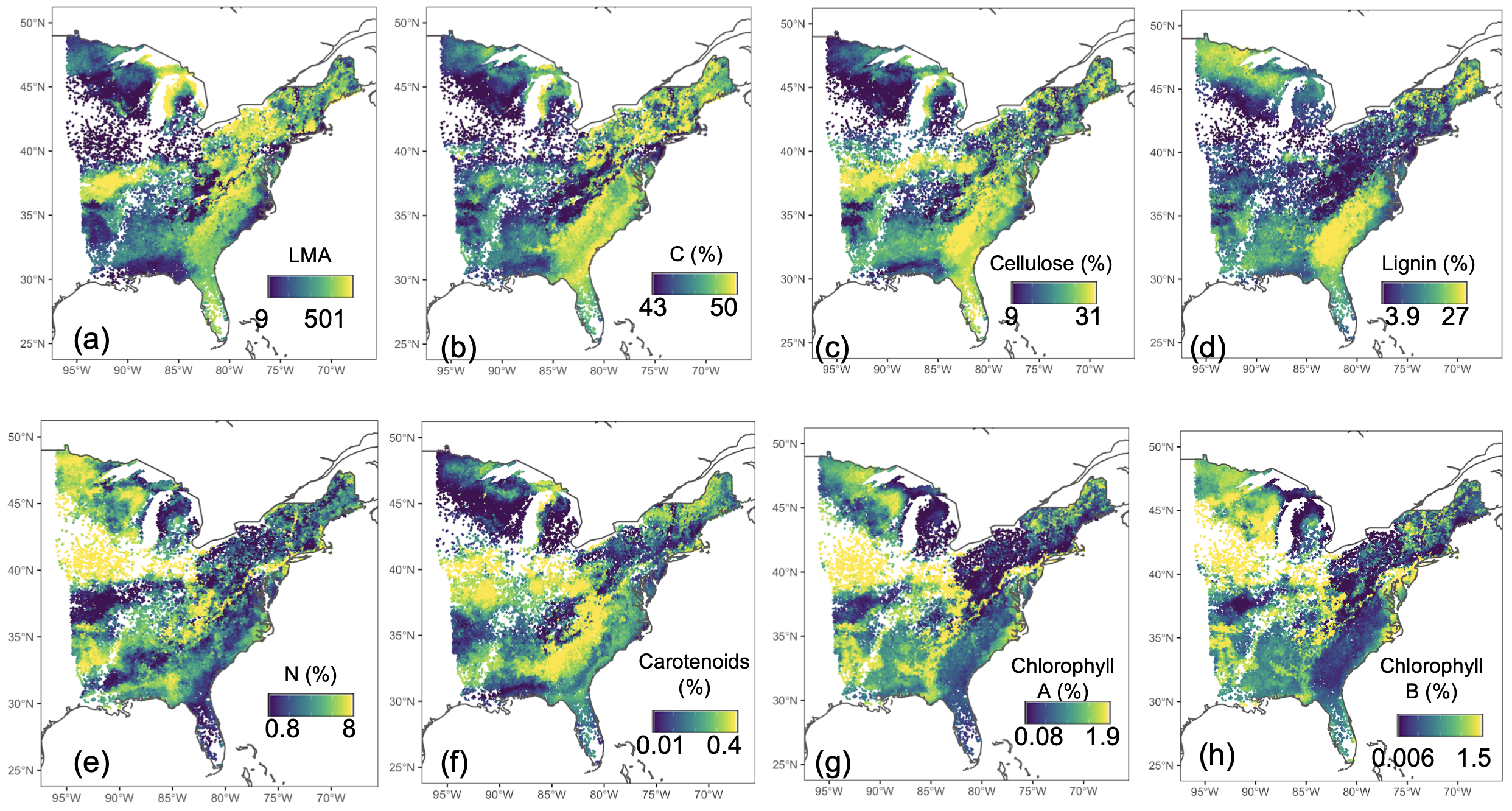

Figure S.13 (continued). Distribution of predictions from the environment-only model for the 8 leaf traits used in this study. Predictions were estimated for each individual tree in the FIA dataset and are shown here averaged across trees within FIA plots. Scale is from low (blue) to high (yellow). From top left to bottom right: chlorophyll A, chlorophyll B, carotenoids, N, C, lignin, cellulose, and LMA. Data to replicate and explore these patterns can be downloaded at https://doi.org/10.5281/zenodo.4647559.

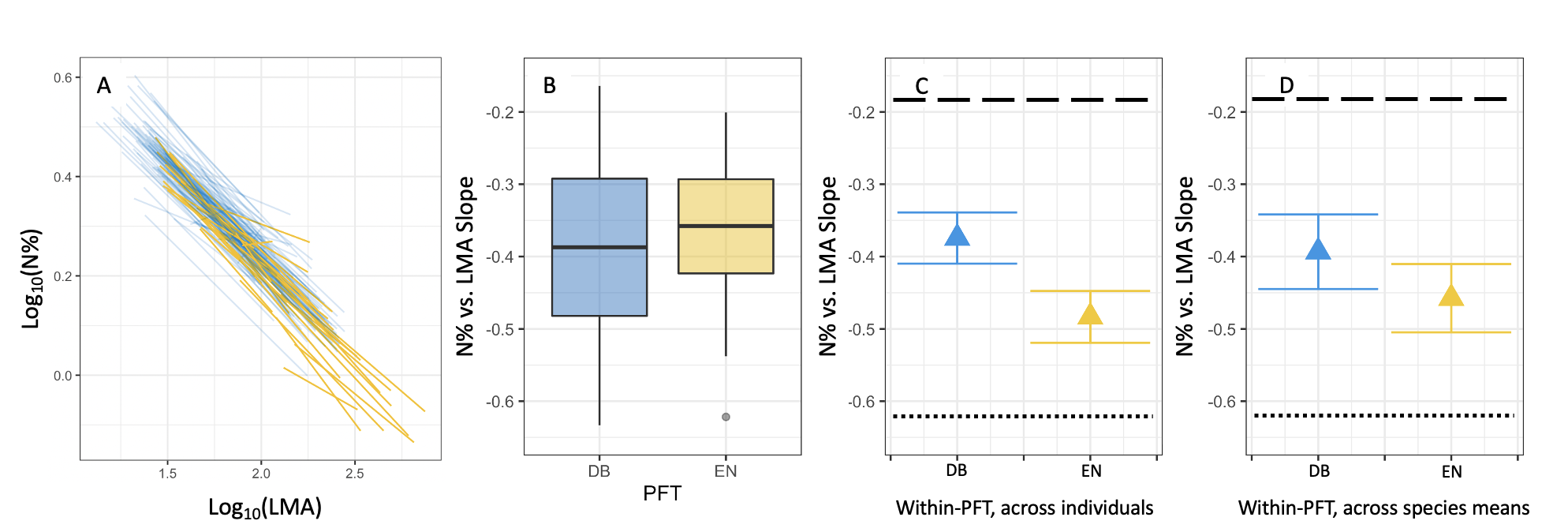

Figure S.14. Relationships between leaf nitrogen percent mass (N%) and leaf mass per area (LMA) across the eastern USA, after rarifying all species abundances to 10 individuals. In this analysis of rarified data, the PFT-level slopes across individuals (C) and across species (D) are similar to each other, suggesting that the differences in these slopes in Fig. 6 are due to differences in species abundance, with commoner species exerting stronger influence on the across-individuals slope. Figure panels and symbols as in Fig. 6. To rarify species abundances, we took a random stratified sample as follows: for each species, we determined the latitudes delimiting each 10% increment of the total number of individuals sampled in FIA plots, and we randomly selected one individual from each of these 10 latitudinal bins.
